# Homoharringtonine Promotes FTO Degradation to Suppress LILRB4-Mediated Immune Evasion in Acute Monocytic Leukemia

**DOI:** 10.1101/2025.03.15.643466

**Authors:** Fangfang Huang, Xiang Luo, Mengyu Zhang, Le Jin, Wenxin Sun, Peihan Chen, Xiuli Hong, Chenyu Xu, Meizhi Jiang, Die Hu, Bin Zhang, Shengwei Hu, Chuanjiang Yang, Rui Gao, Jinzhang Zeng, Quanyi Lu, Qiang Luo, Jun Wu, Siming Chen

## Abstract

Acute monocytic leukemia (AML-M5), a subtype of acute myeloid leukemia, is a highly aggressive malignancy characterized by a poor prognosis, primarily due to the ability of leukemic cells to evade immune surveillance. In this study, we demonstrate that homoharringtonine (HHT), an FDA-approved therapeutic agent for chronic myeloid leukemia (CML), inhibits this immune evasion by targeting the FTO/m6A/LILRB4 signaling pathway in monocytic AML. Utilizing RNA sequencing (RNA-seq) and various functional assays, we reveal that HHT treatment significantly reduces LILRB4 expression at both the RNA and protein levels, suggesting that the effects of HHT on LILRB4 are distinct from its well-established role as a protein synthesis inhibitor. Mechanistically, HHT treatment markedly increases global levels of RNA m6A in THP-1 cells by promoting the degradation of FTO, which subsequently diminishes the expression of its downstream targets, MLL1 and LILRB4. Furthermore, in vitro and in vivo analyses employing monocytic AML cell lines, mouse-derived AML xenograft models, and patient samples collectively support the conclusion that HHT suppresses immune evasion in monocytic AML by reducing LILRB4 expression. Importantly, the downregulation of LILRB4 resulting from HHT treatment enhances the susceptibility of THP-1 cells to CD8^+^ T cell cytotoxicity, accompanied by increased markers of immune activation. Overall, our findings position HHT as a promising clinical agent for enhancing CD8^+^ T cell-based cancer immunotherapy by mitigating immune evasion in monocytic AML.

**Graphical abstract:** 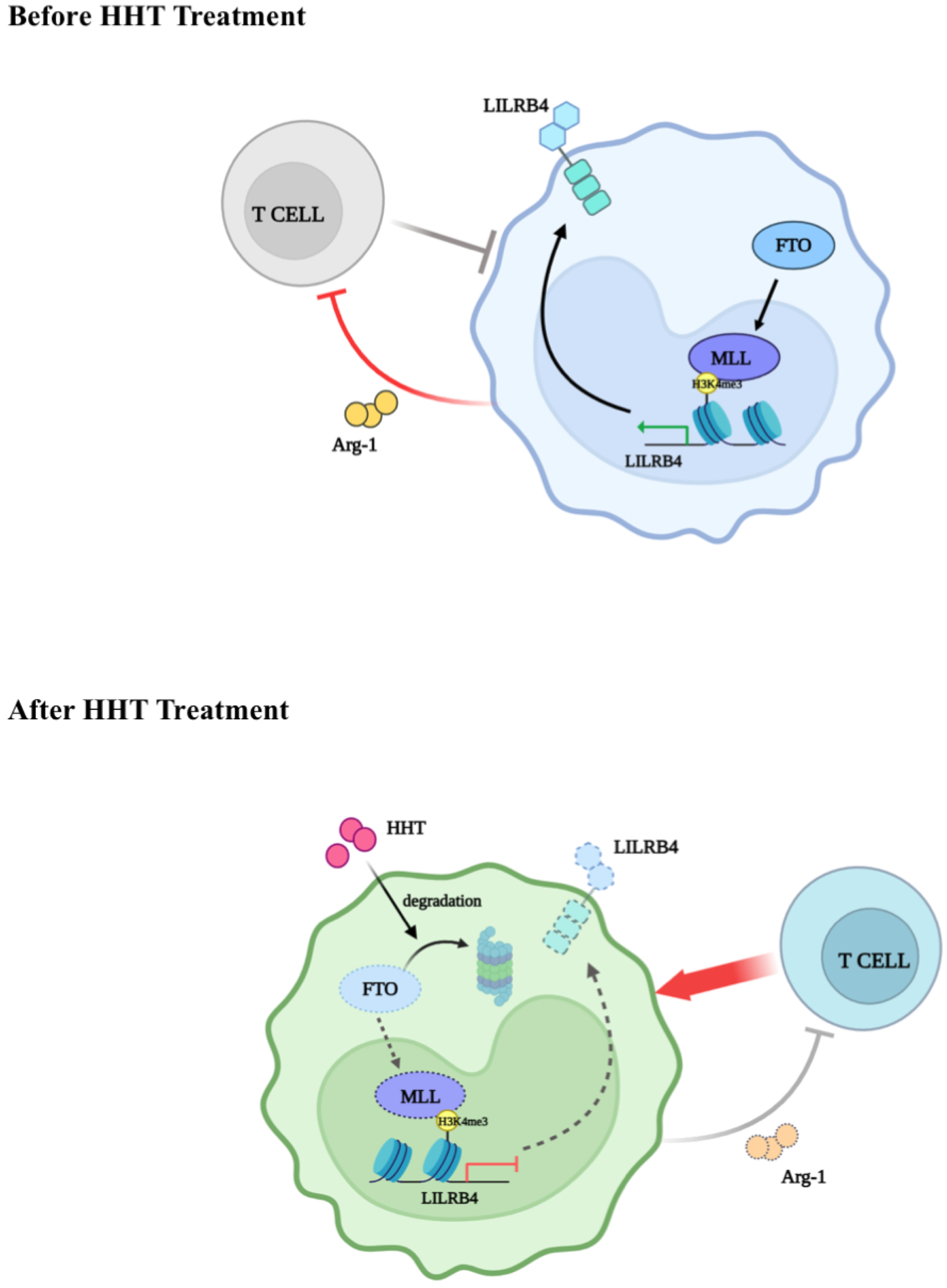

## Introduction

Acute monocytic leukemia (AML-M5), a subtype of acute myeloid leukemia (AML), is characterized by its aggressive behavior and poor prognosis^1–3^. A major challenge in treating AML is its ability to evade immune surveillance, which enables malignant cells to proliferate uncontrollably^4–6^. This immune evasion occurs through various mechanisms, one of which involves the upregulation of programmed death-ligand 1 (PD-L1) on AML cells. PD-L1 interacts with the programmed cell death protein 1 (PD-1) receptor on T cells, inhibiting their cytotoxic responses and facilitating the survival and proliferation of leukemia cells^7–9^. Moreover, AML cells can promote the expansion of myeloid-derived suppressor cells (MDSCs) and regulatory T cells (Tregs), which secrete immunosuppressive molecules such as indoleamine 2,3-dioxygenase (IDO), arginase, and various cytokines^7,10,11^. Collectively, these mechanisms enhance the immune evasion of leukemia cells. A comprehensive understanding of the pathways involved in immune evasion is essential for developing more effective immunotherapeutic strategies for AML, ultimately improving patient outcomes in this challenging disease.

Leukocyte immunoglobulin-like receptor B4 (LILRB4) is an immune inhibitory receptor that is restrictively expressed on monocytic cells and monocytic AML cells^12–14^. Previous studies have shown that LILRB4 suppresses T-cell activity, facilitating the infiltration of leukemia cells into tissues and promoting tumor growth. Inhibiting LILRB4 can reverse this suppression of T-cell activity and impede AML development^12,15,16^. Notably, LILRB4 levels on monocytic AML cells are significantly higher than those on normal monocytes, making it an attractive target for therapeutic intervention^12^. Recent advancements in LILRB4-targeted antitumor research include antibody-drug conjugates (ADCs) that specifically target LILRB4, demonstrating cytotoxicity against monocytic AML cells while exhibiting minimal toxicity to healthy hematopoietic progenitors^17–19^. Additionally, LBL-043, a novel bispecific LILRB4/CD3 antibody, effectively redirects T cells to eliminate AML cells by simultaneously engaging LILRB4 on tumor cells and CD3 on T cells^20–22^. Furthermore, the anti-LILRB4 antibody IO-202 is currently undergoing a Phase 1 clinical trial in combination with azacitidine, targeting patients with AML and chronic myelomonocytic leukemia (CMML)^23,24^. Despite significant advancements in the development of LILRB4-targeting antibodies for treating monocytic AML, research on small molecule compounds that inhibit LILRB4 remains limited^14,25^. This highlights the urgent need for further studies to explore and validate the potential of these compounds in enhancing the efficacy of LILRB4-targeted immunotherapies.

Homoharringtonine (HHT) is a natural alkaloid derived from the plant *Cephalotoxus fortunei.* and is an FDA-approved drug for the treatment of chronic myeloid leukemia (CML)^26–29^. In addition to its use in CML, HHT has been extensively employed in China for treating acute myeloid leukemia (AML) and myelodysplastic syndrome (MDS) ^30–32^. Moreover, HHT demonstrates significant inhibitory effects against various solid tumors, including lung, liver, colorectal, and breast cancers^33–35^. Mechanistically, the anti-tumor properties of HHT are attributed to its binding to the A site of the ribosome, which inhibits the binding of aminoacyl-tRNA during mRNA translation, thereby impeding protein synthesis^33,35,36^. This mechanism of action renders HHT particularly effective against rapidly dividing leukemic cells. Clinical trials and studies have substantiated its efficacy in inducing remission and enhancing patient outcomes, whether administered alone or in combination with other chemotherapeutic agents. However, to our knowledge, the immunotherapeutic potential of HHT in monocytic AML remains unclear, and the underlying molecular mechanisms have yet to be investigated.

In this study, we elucidate that HHT can effectively suppress immune evasion in monocytic AML by inhibiting the FTO/m6A/LILRB4 signaling pathway. Our results demonstrate that HHT enhances intracellular global RNA m6A levels by promoting the degradation of FTO. This, in turn, reduces the expression of its downstream targets, MLL1 and LILRB4. Upon HHT treatment, LILRB4 levels are significantly decreased, which further modulates downstream signaling pathways, including phospho-SHP-2 (Tyr580) and arginase-1 (ARG1). These pathways are crucial for the infiltration of AML cells into tissues and the suppression of T-cell activity^12^. The HHT-mediated downregulation of LILRB4, which contributes to the suppression of immune evasion, was also confirmed in mouse-derived AML xenograft models and patient samples. Collectively, our findings reveal a previously unrecognized mechanism by which HHT-induced reduction in LILRB4 expression can overcome LILRB4-mediated immune escape in monocytic AML. This underscores the significant potential of HHT-based therapies for immunotherapeutic applications.

## Results

### HHT Influences Immune Signaling Pathways in Acute Monocytic Leukemia

To uncover novel mechanisms underlying the therapeutic effects of HHT on monocytic AML, we performed RNA sequencing (RNA-seq) analysis on THP-1 cells treated with either 40 nM HHT or an equivalent volume of dimethyl sulfoxide (DMSO) as a control. Our results revealed that HHT treatment influenced the expression of 4,089 genes, comprising 2,240 downregulated and 1,849 upregulated genes associated with diverse biological functions (log2 fold change ≥ 1, adjusted P ≤ 0.05; Fig. 1A and 1B). Upon increasing the threshold to a log2 fold change ≥ 1.25 and an adjusted P-value of ≤ 0.05 for Gene Ontology (GO) functional enrichment analysis, we found that HHT treatment profoundly impacted several key biological processes, including immune system function, the mitogen-activated protein kinase (MAPK) signaling pathway, and mitotic cell cycle transitions (Fig. 1C). A detailed examination of genes involved in the negative regulation of the immune system pathway highlighted that HHT treatment significantly reduced the expression of the immune checkpoint factor LILRB4, which is restrictively expressed on monocytic AML cells (Fig. 1D and 1E). This finding suggests that HHT treatment may play a role in LILRB4-mediated immune evasion in monocytic AML.

**Figure 1.**
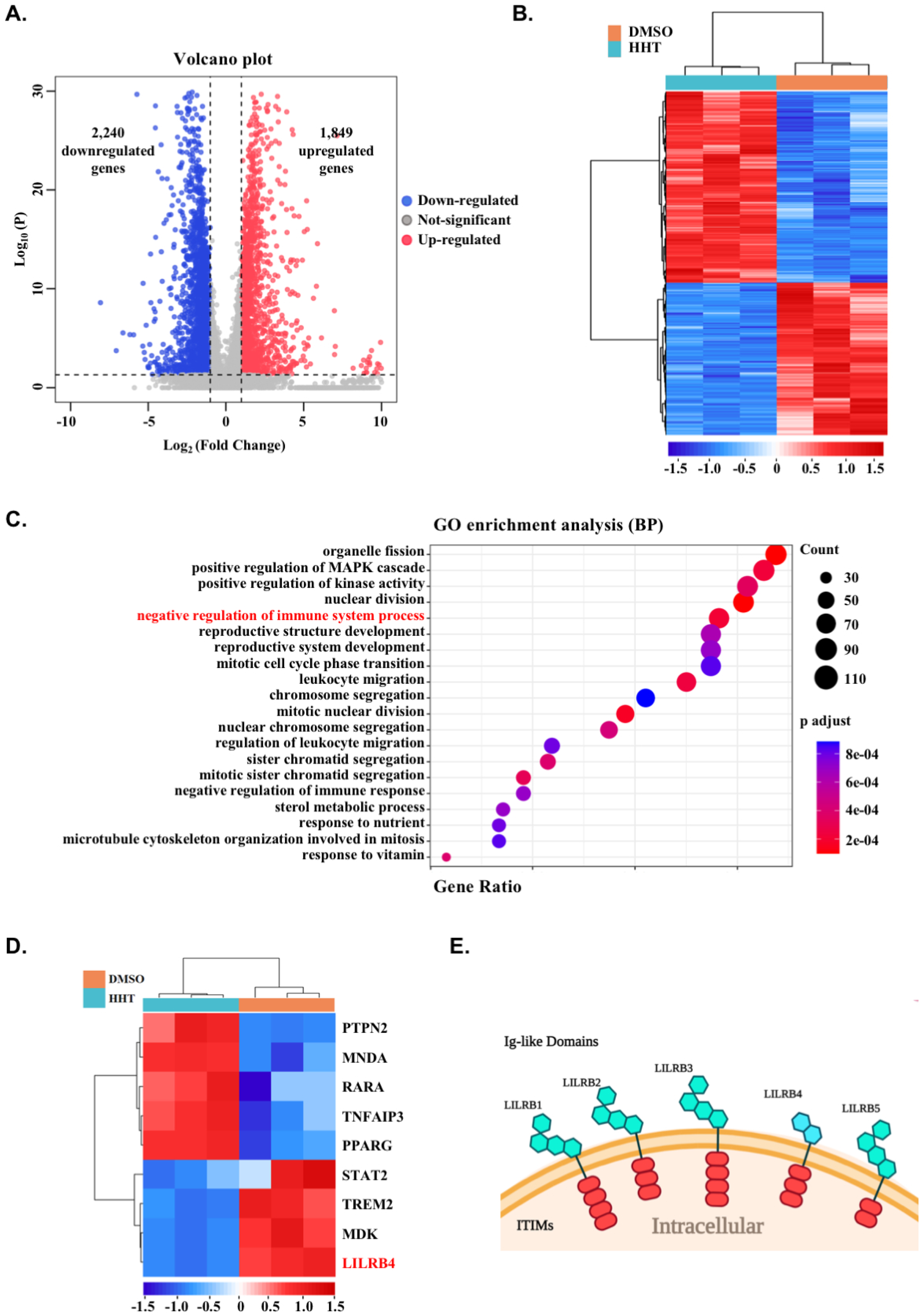
Effects of HHT on Gene Expression and Associated Pathways in THP-1 Cells. (A) Volcano plot illustrating differential gene expression profiles between DMSO (control) and HHT-treated samples. The plot identifies 2,240 downregulated genes (blue) and 1,849 upregulated genes (red), using a threshold of adjusted p-value ≤ 0.05 and log2 fold change ≥ 1. (B) Heatmap visualization of expression patterns for differentially expressed genes in DMSO and HHT treated samples. The clustering dendrogram demonstrates the similarity in gene expression profiles across samples. (C) Gene Ontology (GO) enrichment analysis of biological processes affected by HHT treatment. The bar chart represents the number of genes associated with each biological process, with the most significantly enriched processes related to immune system regulation highlighted. The x-axis indicates the number of genes, while the y-axis lists the biological processes. (D) Immune-Related Pathway Heatmap. This heatmap focuses on the expression changes of genes involved in immune-related pathways, demonstrating the inhibitory effect of HHT on these pathways. (E) Schematic representation of LILRB protein family structure. The diagram illustrates the characteristic structural domains, including extracellular immunoglobulin-like (Ig-like) domains and intracellular immunoreceptor tyrosine-based inhibitory motifs (ITIMs). The schematic was created using BioRender, highlighting the conserved structural features across LILRB family members (LLLRB2, LLLRB3, LLLRB4, LLLRB5).

Among the genes whose expression was altered by HHT treatment, LILRB4, recognized as an immune inhibitory receptor, captured our interest. LILRB family proteins are characterized by their extracellular immunoglobulin-like domains and intracellular immunoreceptor tyrosine-based inhibitory motifs (ITIMs)^12^ (Fig. 1E). Previous studies have demonstrated that LILRB4 interacts with ligands such as apolipoprotein E (APOE) through its extracellular domain, thereby activating the downstream SHP-2-NFκB pathway. This activation leads to the release of arginase-1 (ARG1), which suppresses T cell activity and facilitates immune evasion in monocytic AML^12^. Analysis of the TCGA database via cBioPortal revealed that LILRB4 is restrictively expressed on monocytic AML cells, particularly in the French-American-British M4 and M5 subtypes (FAB M4 and M5 AML subtypes) (Fig. S1A), consistent with previous findings^12^. Furthermore, Kaplan-Meier Plotter analysis indicated that AML patients with high LILRB4 expression exhibit significantly lower survival rates compared to those with low expression (Fig. S1B). These findings highlight the critical role of LILRB4 in influencing survival and immune suppression in patients with AML-M4/M5, establishing it as a promising therapeutic target for treating monocytic AML.

### HHT Decreases LILRB4 Expression at Both Transcriptional and Translational Levels

To further validate the effects of HHT on LILRB4 expression, we selected the acute monocytic leukemia cell line THP-1 and treated it at a density of 2.5 × 10^5^ cells/ml with varying concentrations of HHT (ranging from 0 nM to 160 nM) using a two-fold dilution series over 48 and 72 hours. Reverse transcription-quantitative PCR (RT-qPCR) analyses demonstrated that exposure of THP-1 cells to HHT resulted in a dose-dependent decrease in LILRB4 mRNA expression, consistent with our previous RNA sequencing findings (Fig. 2A-C). To evaluate the impact of HHT on LILRB4 protein expression, we employed two complementary methodologies. First, flow cytometry was used to analyze the changes in LILRB4 protein levels on the surface of THP-1 cells following treatment with various concentrations of HHT and subsequent staining with an LILRB4-specific antibody. Consistent with our RT-qPCR results, flow cytometric analysis indicated a significant reduction in LILRB4 protein expression in THP-1 cells treated with HHT compared to those treated with DMSO control (Fig. 2D-F). Furthermore, the dose-dependent reductions in LILRB4 protein expression were also confirmed through Western blot analysis (Fig. 2G). Collectively, these findings suggest that HHT downregulates LILRB4 expression at both the transcriptional and translational levels.

**Figure 2.**
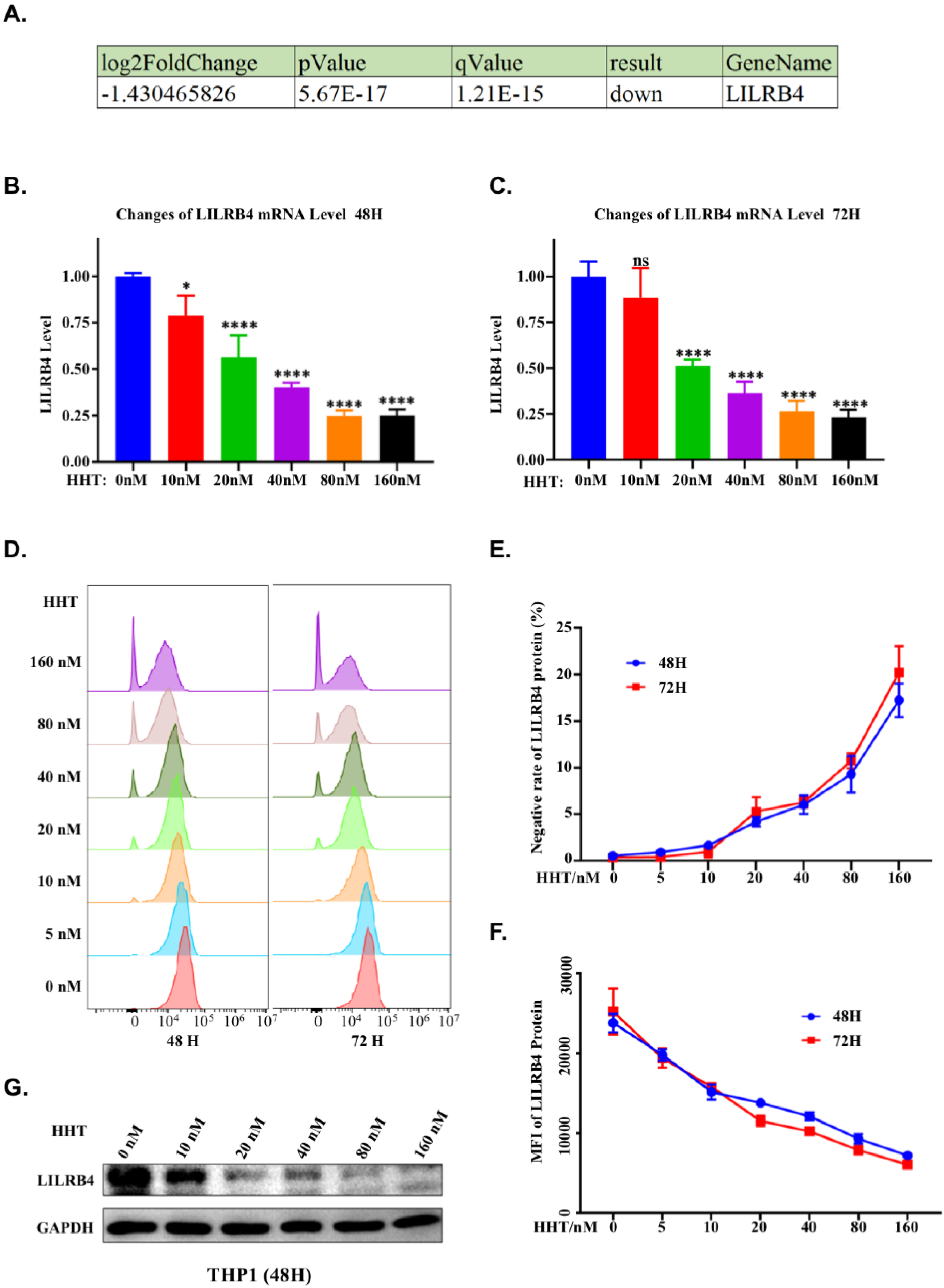
HHT Inhibits LILRB4 Expression at Both Transcriptional and Translational Levels in THP-1 Cells. (A) RNA-seq analysis indicating the downregulation of LILRB4 transcriptional levels, with a log2 fold change of -1.430465826, p-value of 5.67E-17, and q-value of 1.21E-15, confirming significant downregulation. (B-C) Bar graphs showing the mRNA levels of LILRB4 in THP-1 cells treated with a gradient of 160 nM HHT for 48 and 72 hours. The results demonstrate a decrease in LILRB4 mRNA levels over time. (D-F) Flow cytometry histograms depicting LILRB4 expression in THP-1 cells treated with HHT. As the concentration of HHT increases, the mean fluorescence intensity of LILRB4 decreases, and the proportion of LILRB negative cells increases. (G) Western blot analysis revealing a reduction in LILRB4 protein expression after 48 hours of treatment with increasing concentrations of HHT. GAPDH is used as a loading control.

HHT is widely recognized as a protein synthesis inhibitor. However, our RNA sequencing (RNA-seq) and subsequent RT-qPCR results demonstrate that HHT treatment significantly reduces LILRB4 expression at both the protein and mRNA levels. This observation prompts us to consider that the effects of HHT on LILRB4 protein levels may different from its conventional role as a protein synthesis inhibitor. Therefore, in this study, we aim to elucidate the mechanisms by which HHT downregulates LILRB4 protein expression, with a particular focus on the upstream transcriptional regulation of LILRB4 in response to HHT treatment.

### HHT Reduces FTO and MLL1 Complex Levels in Acute Monocytic Leukemia

Epigenetic dysregulation is a hallmark of acute myeloid leukemia (AML), characterized by abnormal epigenetic patterns that lead to aberrant gene expression^37–40^. Given that HHT treatment modulates LILRB4 mRNA levels, we sought to investigate whether HHT regulates LILRB4 protein expression by altering the epigenetic landscape in monocytic AML. Extensive research has shown that LILRB4 protein expression is subject to epigenetic modulation^41–43^. Notably, FTO, an mRNA N6-methyladenosine (m6A) demethylase, has been identified as a key upstream regulator of LILRB4 in AML cells^44^. Specifically, treating AML cells with an FTO inhibitor or knocking down FTO expression results in reduced LILRB4 expression^44^. In contrast, treatment with decitabine, a hypomethylating agent, increases LILRB4 expression, presumably by upregulating FTO^44^. Additionally, an analysis of H3K4me3 chromatin immunoprecipitation sequencing (ChIP-seq) data from the MV4-11 and THP-1 cell lines has revealed significant enrichment of H3K4me3 modification peaks in the promoter regions of the LILRB4 gene (Fig.S2A)^45^. This finding suggests that LILRB4 expression may also be regulated by the MLL1 protein complex, which catalyzes the H3K4me3 modification to activate gene expression. Collectively, these observations indicate that epigenetic regulatory factors, such as FTO and MLL1, likely play pivotal roles in modulating LILRB4 expression in AML cells in response to HHT. Despite these insights, the precise molecular mechanisms by which HHT downregulates LILRB4 expression remain unclear.

To explore the potential epigenetic factors that modulate LILRB4 expression in response to HHT treatment, we examined the impact of HHT on various epigenetic regulators. In acute myeloid leukemia (AML), dysregulation of N6-methyladenosine (m6A) modification is recognized as a primary mechanism underlying tumor growth and immune evasion through its effects on gene expression^46–48^. This regulatory network comprises three key components: the “Writer” complex, which includes METTL3, METTL14, and WTAP and facilitates the addition of m6A marks to RNA adenosines; “Reader” proteins, such as IGF2BP1/2, which engage with m6A-modified RNA to influence RNA stability and translational efficiency; and “Eraser” proteins, such as FTO and ALKBH5, which remove m6A marks, thereby counteracting these modifications. This system intricately regulates RNA functionality, with m6A playing a crucial role in gene expression and immune evasion in AML (Fig. 3A)^47^. Additionally, chromosomal translocations frequently occur in AML, leading to the fusion of MLL with other genes, such as AF4, AF9, or ENL, which is another significant feature of AML^49^. In normal cells, the MLL gene consists of two distinct domains: the N-terminal domain (MLL-N) and the C-terminal domain (MLL-C). The formation of MLL fusion proteins can result in abnormal gene expression and chromatin remodeling, thereby promoting tumor growth in AML (Fig. 3B)^50^. Expanding our investigation beyond the previously mentioned proteins and complexes implicated in AML, we further explored the responses of additional epigenetic factors or complexes to HHT treatment. These include the Polycomb Repressive Complex 2 (PRC2), which catalyzes the trimethylation of histone H3 at lysine 27, and the protein arginine methyltransferases (PRMTs), responsible for the dimethylation of histone H4 at arginine 3. To this end, we treated THP-1, MV4-11, and NOMO-1 cells with varying concentrations of HHT for 48 hours and observed a significant decrease in the protein levels of FTO, DOT1L, and MLL1 across all three cell lines following HHT treatment (Fig. 3C-D and S3A). Notably, unlike FTO, the m6A writers and readers, including METTL3, METTL14, and IGF2BP2, demonstrated insensitivity to HHT treatment under the same experimental conditions. Given that FTO and MLL1 have been proposed as upstream regulators of LILRB4, we questioned whether HHT downregulates LILRB4 expression by reducing the levels of MLL1 and FTO.

**Figure 3.**
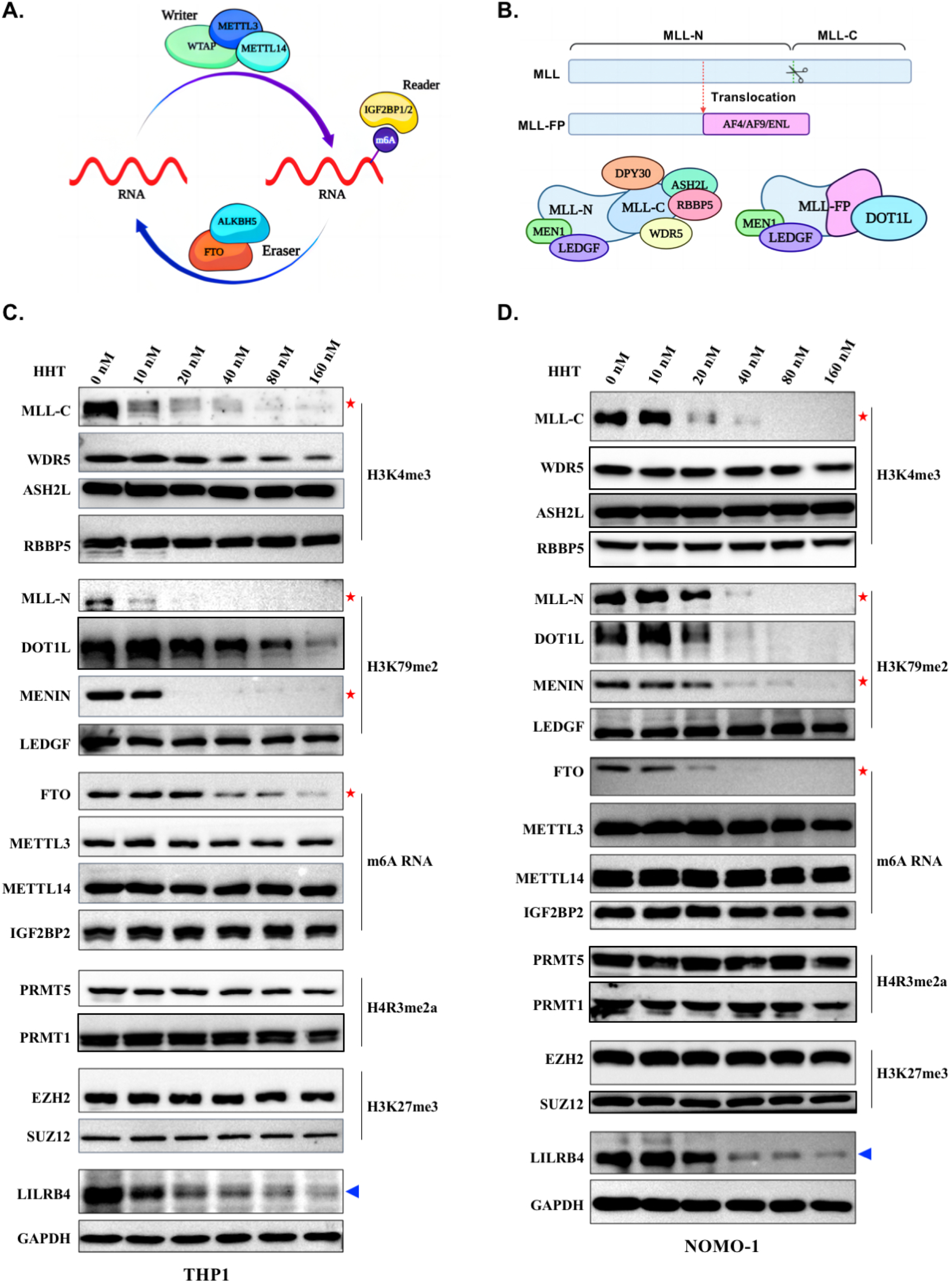
HHT Regulates the Expression of Epigenetic Regulatory Factors. (A) This panel illustrates the m6A RNA methylation complex, delineating the roles of writer proteins (e.g., METTL3, METTL14, WTAP), eraser proteins (e.g., ALKBH5, FTO), and reader proteins (e.g., IGF2BP1/2) in shaping RNA methylation landscapes. The schematic was rendered using BioRender.com. (A) The upper diagram illustrates the domain structure of wild-type MLL and MLL Fusion Proteins (MLL-FPs). Wild-type MLL is cleaved by taspase1 into MLL-N and MLL-C domains. Red arrows indicate frequent translocation breakpoints in AML, leading to the formation of common MLL-FPs (MLL-AF4, MLL-AF9, MLL-ENL). These translocations result in chimeric proteins that disrupt MLL function, contributing to leukemogenesis through dysregulated gene expression and epigenetic modifications. The lower diagrams show: on the left, the MLL complex components (MLL-N, MLL-C, DPY30, ASH2L, RBBP5, WDR5) involved in H3K4 methylation; on the right, MLL-FPs associated with DOT1L, MEN1, and LEDGF, highlighting their role in altered gene regulation. Created with BioRender.com. (C-D) Western blot analysis was performed to evaluate the expression of epigenetic complexes in THP-1 and NOMO-1 cells following treatment with a range of 0-160 nM HHT for 48 hours. The analysis targeted components of the MLL and MLL-FPs complex (MENIN, MLL-N, MLL-C), the H3K27me3 complex (EZH2, SUZ12), the m6A complex (METTL3, METTL14, FTO, IGF2BP2), and H4R3me2a-associated proteins (PRMT1, PRMT5). The findings reveal a marked reduction in the expression of the m6A demethylase FTO and pivotal components of the MLL and MLL-FPs complex, namely MENIN, MLL-N, and MLL-C, in response to HHT treatment.

### HHT Decreases LILRB4 Expression by Reducing FTO Levels

To investigate the potential correlation between FTO, MLL1, and LILRB4, we generated stable FTO knockdown and overexpression cell lines using human monocytic acute myeloid leukemia (AML) cell lines, including THP-1, NOMO-1, and MV4-11. The efficiency of FTO knockdown and overexpression was confirmed by RT-qPCR and Western blotting, respectively (Fig. 4A and 4C). Our results demonstrated that FTO knockdown significantly decreased LILRB4 levels at both the mRNA and protein levels. Conversely, overexpression of wild-type FTO, but not its enzymatically inactive mutant, robustly enhanced LILRB4 expression (Fig. 4B and 4C). Consistent findings were observed in NOMO-1 and MV4-11 cell lines, where LILRB4 levels were tightly correlated with FTO levels (Fig. 4D-F and Fig. S4A-C). Collectively, these data highlight the critical role of FTO in regulating LILRB4 expression in human monocytic AML cell lines.

**Figure 4.**
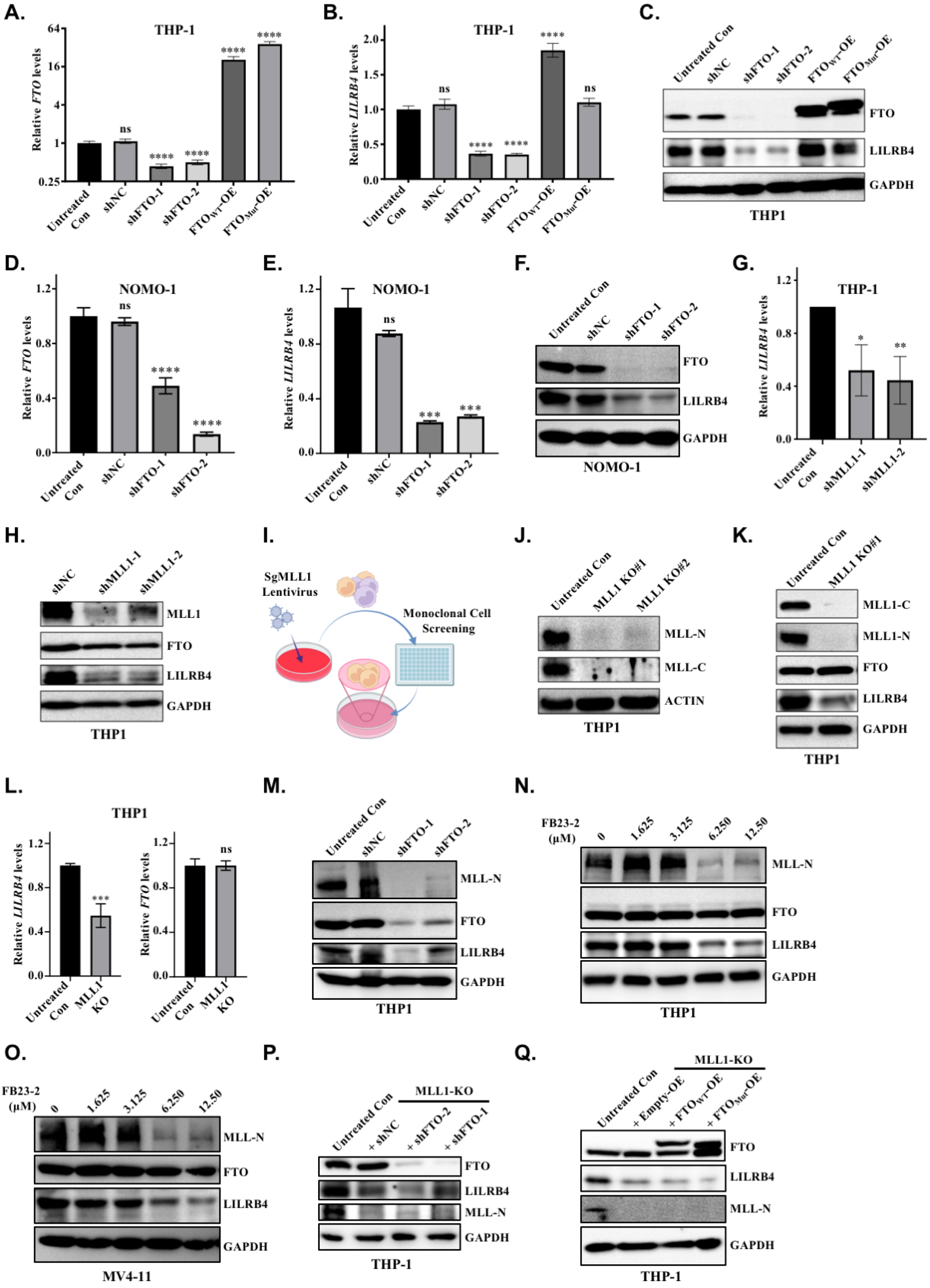
FTO Regulates LILRB4 Expression in an MLL-Dependent Manner. (A-B) RT-qPCR analysis reveals that knockdown of FTO results in decreased LILRB4 mRNA expression, while overexpression of wild-type FTO (FTO^WT^), but not its mutant form (FTO^Mut^), enhances LILRB4 transcription levels. (A) Western blot analysis of THP-1 cells with FTO knockdown and overexpression demonstrates that FTO positively regulates LILRB4 protein expression. (D-E) RT-qPCR analysis of NOMO-1 cells confirms that FTO knockdown downregulates LILRB4 mRNA expression. (F) Western blot analysis corroborates that the knockdown of FTO in NOMO-1 cells reduces LILRB4 protein expression. (G) RT-qPCR analysis indicates that MLL knockdown in THP-1 cells decreases LILRB4 mRNA levels. (H) Western blot analysis shows that MLL knockdown in THP-1 cells downregulates LILRB4 protein expression without affecting FTO protein levels. (I) A schematic representation of the monoclonal cell line screening process for MLL knockout in THP-1 cells is depicted, generated using BioRender. (J-L) Western blot and RT-qPCR analyses of MLL knockout cell lines reveal that MLL knockout does not affect FTO mRNA and protein expression but downregulates LILRB4 mRNA and protein expression. (M) Western blot analysis of FTO knockdown cells demonstrates a decrease in MLL expression. (N-O) Treatment of THP-1 and MV4-11 cells with the FTO enzyme inhibitor FB23-2 leads to reduced expression of MLL and LILRB4, exhibiting a phenotype similar to that observed in FTO knockdown cells. (P-Q) Further knockdown or overexpression of FTO in MLL knockout cell lines shows no significant effect on LILRB4 expression, suggesting that FTO’s regulatory role on LILRB4 is MLL-dependent.

We treated human monocytic acute myeloid leukemia (AML) cell lines, including THP-1, NOMO-1, and MV4-11, with HHT and observed a significant reduction in MLL1 levels (Fig. 3C-D and Fig. S3A). Additionally, analysis of publicly available H3K4me3 chromatin immunoprecipitation sequencing (ChIP-seq) datasets revealed an enrichment of H3K4me3 modifications at the LILRB4 promoter in monocytic AML cells (Fig. S2A). Given that MLL1 is a key enzyme responsible for H3K4me3 modifications, these observations suggest that MLL1 may directly regulate LILRB4 expression by altering the H3K4me3 modification on the LILRB4 promoter in response to HHT treatment. To elucidate the regulatory role of MLL1 in LILRB4 expression, we generated stable THP-1 cell lines with MLL1 knockdown and assessed LILRB4 expression levels using Western blot and RT-qPCR analyses. Our results demonstrated a significant decrease in LILRB4 expression at both the protein and mRNA levels following MLL1 knockdown (Fig. 4G-H). These findings indicate that MLL1 regulates LILRB4 expression at both the transcriptional and translational levels, suggesting that MLL1-mediated epigenetic modifications play a significant role in the regulation of LILRB4 expression in response to HHT treatment.

To clarify the hierarchical relationship between FTO and MLL1 in regulating LILRB4 expression, we employed CRISPR-mediated genome editing to knockout endogenous MLL1 in THP-1 cells (Fig. 4I-J). Subsequent RT-qPCR and Western blot analyses revealed that MLL1 knockout significantly downregulated LILRB4 levels at both the RNA and protein levels, while FTO levels remained unchanged (Fig. 4K-L). This finding is consistent with our previous results from MLL1 knockdown cell lines (Fig. 4G-H), indicating that MLL1 acts downstream of FTO to specifically regulate LILRB4 expression. Conversely, when FTO was knocked down in THP-1 cells, expression levels of both MLL1 and LILRB4 were significantly reduced (Fig. 4M). This suggests that FTO acts upstream of MLL1 to modulate LILRB4 expression. Furthermore, treatment of monocytic AML cell lines, including THP-1, MV4-11, and NOMO-1, with the FTO enzymatic inhibitor FB23-2 resulted in a significant dose-dependent decrease in the protein levels of both LILRB4 and MLL1 (Fig. 4N-O and Fig. S4D). This confirms that FTO regulates LILRB4 and MLL1 expression in an enzymatic activity-dependent manner. Finally, stable knockdown or overexpression of FTO in MLL1 knockout (MLL1-KO) THP-1 cells failed to rescue LILRB4 expression (Fig. 4P-Q). This further implies that FTO’s regulation of LILRB4 is highly dependent on MLL1. Collectively, these findings demonstrate that FTO acts upstream of MLL1 to regulate LILRB4 expression in acute monocytic leukemia, and this regulatory process is mediated through FTO’s mRNA N6-methyladenosine (m6A) demethylase activity.

### HHT Inhibits GSK3β Phosphorylation to Facilitate FTO Degradation

The experimental results presented above demonstrate that the HHT drug downregulates FTO protein levels, which subsequently leads to a reduction in LILRB4 expression in monocytic AML cells. To elucidate the mechanism underlying this HHT-mediated decrease in FTO protein levels, we first examined FTO mRNA levels in THP-1 cells treated with a range of HHT concentrations from 0 to 160 nM using RT-qPCR. The results revealed that HHT does not downregulate FTO mRNA levels but instead slightly upregulates them, consistent with our previous RNA sequencing results (Fig. 5A and Fig. S5A). This suggests that the observed decrease in FTO protein levels is likely occurring at the translational or post-translational level rather than at the transcriptional level. Given that HHT is known to exert its biological effects by inhibiting protein synthesis, we investigated whether the reduction in FTO protein levels is due to HHT promoting FTO protein degradation or inhibiting its synthesis. To address this question, we used cycloheximide (CHX), a well-known protein synthesis inhibitor, to block the synthesis of new FTO protein in THP-1 cells. This approach allowed us to specifically observe the effects of HHT on the degradation of existing FTO protein. To this end, we treated THP-1 cells with CHX either alone or in combination with HHT. Our findings indicated that the rate of FTO protein degradation in cells treated with both CHX and HHT was significantly greater than that in cells treated with CHX alone (Fig. 5B-C). This result suggests that HHT likely downregulates FTO protein levels by promoting the degradation of FTO protein, rather than by inhibiting its synthesis.

**Figure 5.**
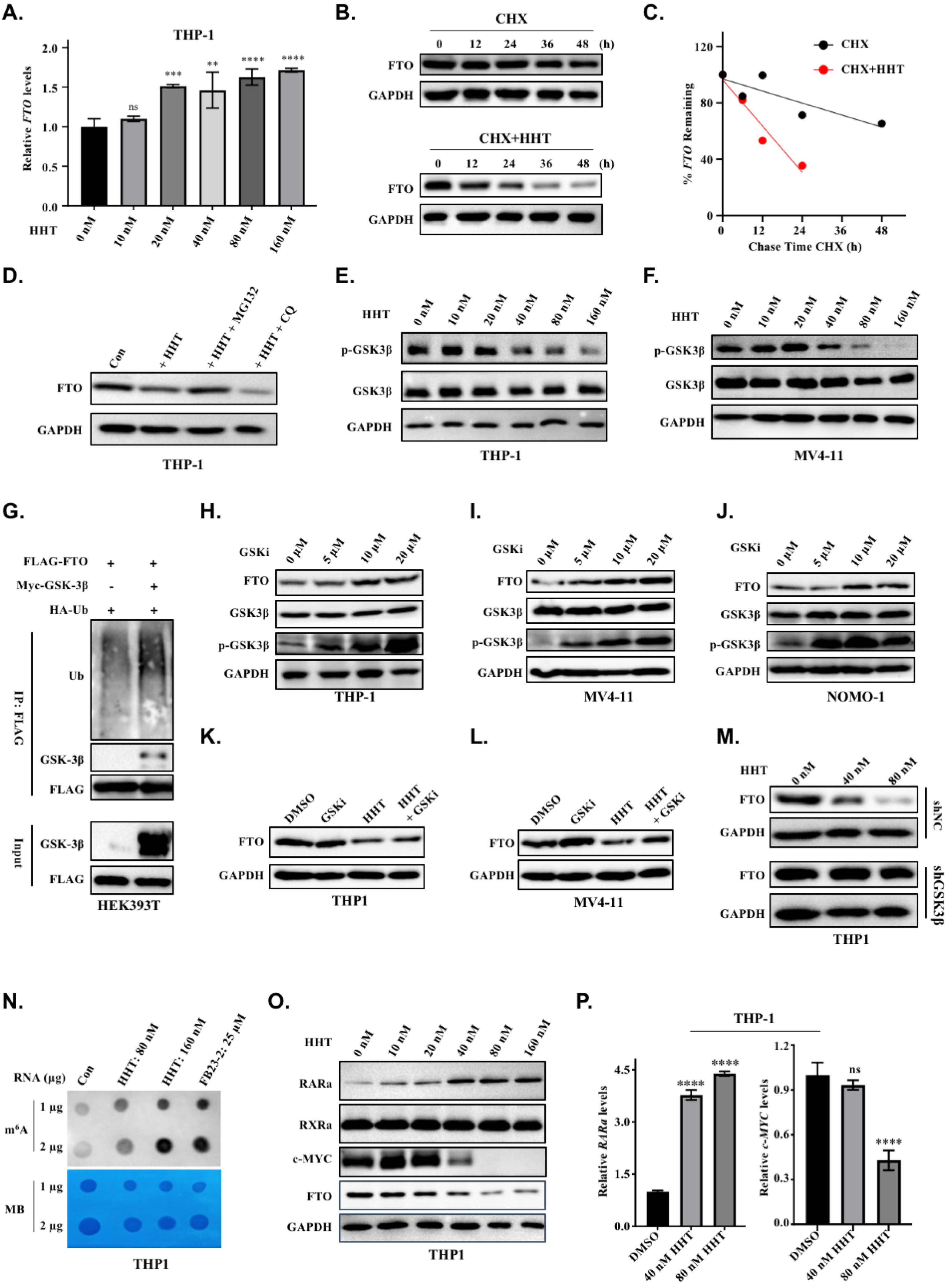
HHT Promotes FTO Degradation by Suppressing GSK3β Phosphorylation Activity. (A) RT-qPCR analysis shows that treatment with HHT slightly upregulates FTO expression at the mRNA level in THP-1 cells. (B-C) Cycloheximide chase assay demonstrates accelerated FTO protein degradation upon HHT co-treatment in THP-1 cells. Protein synthesis inhibition by cycloheximide alone or in combination with HHT reveals that HHT enhances FTO protein turnover. (D) Inhibition of proteasomal degradation by MG132, but not lysosomal inhibition by chloroquine (CQ), rescues HHT-induced FTO downregulation, indicating that FTO degradation occurs primarily through the proteasome pathway. (E-F) Dose-dependent reduction of GSK3β phosphorylation at Ser9 (P-GSK3β) in THP-1 and MV4-11 cells following HHT treatment (0-160 nM), as demonstrated by western blot analysis, indicating HHT-mediated activation of GSK3β signaling. (G) Co-immunoprecipitation analysis in HEK 293T cells demonstrates a physical interaction between GSK3β and FTO, with GSK3β promoting FTO ubiquitination and subsequent degradation. (H-J) Pharmacological inhibition of GSK3β kinase activity in THP-1, MV4-11, and NOMO-1 cells leads to elevated FTO protein levels, demonstrating that GSK3β negatively regulates FTO stability. (K-L) GSK3β kinase inhibitor treatment rescues HHT-mediated FTO degradation in THP-1 and MV4-11 cells, demonstrating that HHT-induced FTO downregulation depends on GSK3β activity. (M) GSK3β knockdown cells exhibit resistance to HHT-induced FTO degradation compared to control cells, demonstrating that GSK3β is required for HHT-mediated downregulation of FTO. (N) Dot blot analysis indicates that HHT treatment enhances global m6A methylation, mimicking the effect of the FTO inhibitor FB23-2. (O-P) Western blot and RT-qPCR analyses reveal that HHT treatment modulates the expression of FTO downstream targets, including RARA and c-MYC, at both the protein and mRNA levels.

Cellular protein homeostasis is maintained through two primary protein degradation systems: the ubiquitin-proteasome system (UPS) and the autophagy-lysosome pathway (ALP), both of which are responsible for the degradation of most cellular proteins^51^. To determine which pathway is involved in HHT-mediated FTO protein degradation, we treated THP-1 cells with either 5 mM MG132 (a proteasome inhibitor) or 25 μM chloroquine (an autophagy inhibitor) for 2 hours, followed by HHT treatment for an additional 36 hours. Our results showed that MG132 inhibited the ability of HHT to induce FTO protein degradation, whereas chloroquine had no effect on HHT-induced FTO degradation (Fig. 5D). These findings confirm that the proteasome pathway, but not the autophagy pathway, is responsible for HHT-mediated FTO protein degradation.

Previous studies have demonstrated that GSK3β facilitates the phosphorylation of FTO, thereby enhancing its ubiquitin-mediated degradation in both mouse embryonic stem cells and human cervical cancer cells^52,53^. To determine whether GSK3β is involved in HHT-mediated FTO degradation in THP-1 cells, we performed Western blot analysis to examine the phosphorylation status of GSK3β at Ser9, as well as its total protein levels, following treatment with various concentrations of HHT (0–160 nM). The results revealed a significant decrease in GSK3β phosphorylation at Ser9, while total GSK3β levels remained unchanged (Fig. 5E). Similar findings were observed in MV4-11 cells (Fig. 5F). These observations imply that HHT induces FTO degradation by inhibiting GSK3β phosphorylation at Ser9, thereby activating GSK3β and promoting FTO degradation. To further elucidate the role of GSK3β in FTO degradation via polyubiquitination, we conducted co-transfection experiments in HEK293T cells using FLAG-tagged FTO and HA-tagged ubiquitin, with or without GSK3β. We then immunoprecipitated FTO using an anti-FLAG antibody and performed Western blotting for ubiquitin using an anti-HA antibody. Our results demonstrated that GSK3β binds to FTO and enhances its polyubiquitination compared to HEK293T cells without GSK3β co-transfection (Fig. 5G). We further treated THP-1 cells with the HHT drug and performed immunoprecipitation to assess the ubiquitination levels of endogenous FTO protein. The experimental results demonstrated that HHT treatment led to an increase in the ubiquitination of endogenous FTO protein within the cells (Fig. S5B). Collectively, these data indicate that HHT induces FTO degradation by inhibiting GSK3β phosphorylation, thereby activating GSK3β and promoting FTO ubiquitination and subsequent degradation.

We next utilized the GSK3β inhibitor (SB 415286) to modulate GSK3β phosphorylation in THP-1 cells and subsequently examined its impact on FTO protein levels. Western blot analysis revealed that treatment with SB 415286 increased GSK3β phosphorylation at Ser9, which was associated with elevated FTO protein levels in monocytic AML cells, including THP-1, NOMO-1, and MV4-11 cells (Fig. 5H-J). These findings were consistent with our observations in monocytic AML cells treated with HHT, where HHT treatment resulted in GSK3β dephosphorylation and a corresponding decrease in FTO protein levels. Furthermore, the GSK3β inhibitor effectively rescued HHT-mediated FTO degradation, highlighting that HHT-induced FTO degradation is highly dependent on GSK3β activity (Fig. 5K-L). Notably, knocking down GSK3β using shRNA in THP-1 cells also blocked HHT-induced FTO protein degradation (Fig. 5M). Collectively, these results demonstrate that HHT treatment induces GSK3β dephosphorylation and activation, which subsequently drives FTO degradation in monocytic AML cells.

### HHT Promotes FTO Degradation to Upregulate Global m6A Levels and Modulate Downstream Targets

As an m6A RNA demethylase, FTO regulates cellular m6A modification levels. Given that HHT induces FTO degradation, we investigated whether HHT treatment alters global m6A modification levels in monocytic AML cells. To this end, we conducted m6A dot blot assays to assess changes in m6A abundance following HHT treatment. The results revealed a significant increase in m6A abundance in the transcriptome of THP-1 cells treated with HHT, comparable to levels observed after treatment with the FTO inhibitor FB23-2 (Fig. 5N). To rule out the possibility that the increase in m6A levels is due to direct inhibition of FTO demethylase activity by HHT, we performed in vitro enzymatic assays using purified recombinant FTO protein. The results indicated that HHT does not inhibit FTO enzymatic activity (Fig. S5C-D). Collectively, these findings suggest that HHT treatment in monocytic AML cells enhances global m6A modification levels through FTO protein degradation, rather than through direct inhibition of FTO enzymatic activity.

In acute myeloid leukemia (AML) cells, both FTO knockdown and FTO inhibition have been shown to increase the protein levels of RARα while downregulating CEBPA and MYC^54–57^. Consistent with these regulatory effects, our treatment of THP-1 cells with HHT similarly dysregulated these downstream target genes of FTO. Specifically, we observed a dose-dependent degradation of FTO protein across a range of HHT concentrations, which was accompanied by decreased MYC protein levels and increased RARα protein levels. As a control, another nuclear receptor protein, RXRα, remained unchanged (Fig. 5O). Notably, HHT also exhibited regulatory effects on the mRNA levels of RARα and MYC (Fig. 5P). These findings further support the conclusion that HHT modulates the expression of RARα and MYC at both the mRNA and protein levels by targeting FTO. Collectively, these results confirm that HHT exerts its effects through the degradation of FTO protein, thereby altering the expression of its downstream targets.

### HHT Sensitizes Monocytic AML Cells to T Cell Cytotoxicity by Reducing LILRB4

LILRB4, an immune inhibitory receptor predominantly expressed on monocytic acute myeloid leukemia (AML) cells, is known to suppress T cell activity^12^. In this study, we demonstrate that HHT reduces LILRB4 levels by promoting the degradation of FTO. To investigate whether HHT can sensitize monocytic AML cells to T cell cytotoxicity through the reduction of LILRB4 levels, we generated a stable GFP-expressing THP-1 cell line using a lentiviral system and isolated T cells from the peripheral blood of healthy volunteers. We then developed a co-culture assay to measure the cell-mediated cytotoxicity of effector CD8^+^ T cells against THP-1 cells (Fig. 6A). In these assays, THP-1 cells were pre-treated with HHT or DMSO to decrease endogenous LILRB4 levels (Fig. 6B), washed to remove residual HHT, and then co-cultured with T cell. T cells exhibited heightened sensitivity to HHT-pretreated THP-1 cells, resulting in enhanced cytotoxic activity against THP-1 cells (Fig. 6C). These findings suggest that HHT sensitizes monocytic AML cells to T cell cytotoxicity by reducing LILRB4 levels, thereby enhancing the antitumor activity of T cells.

**Figure 6.**
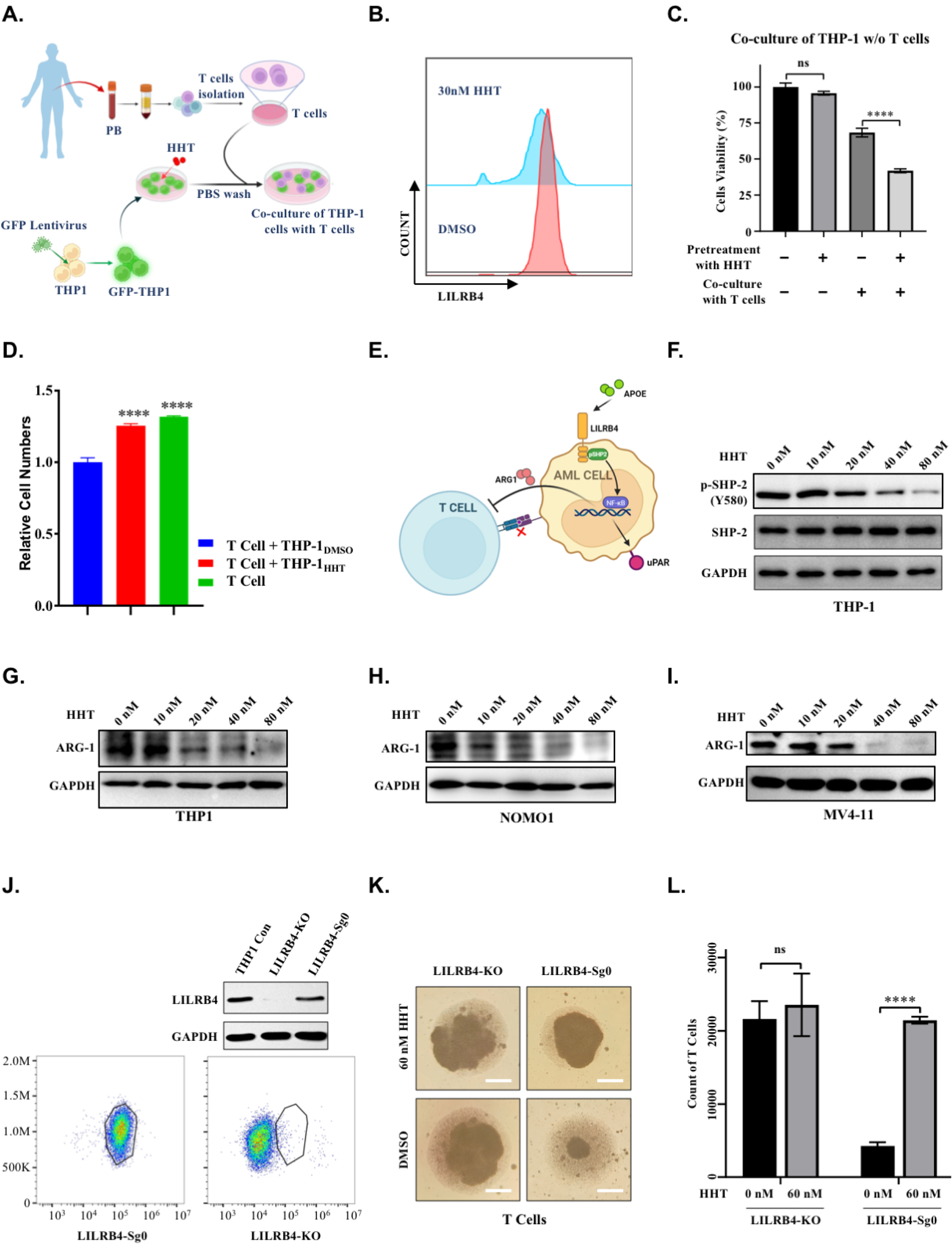
HHT Suppresses Immune Evasion in Acute Myeloid Leukemia (AML) (A) Schematic diagram of the experimental workflow for THP-1-T cell co-culture system, including T cell isolation, GFP labeling through lentiviral transduction, and HHT treatment conditions. Diagram created using BioRender. (B) Flow cytometry analysis demonstrates that 30 nM HHT treatment reduces LILRB4 surface expression on THP-1 cells. (C) THP-1 cells were pre-treated with 30 nM HHT or DMSO control for 48 hours, washed to remove residual compounds, and co-cultured with T cells (1:1 ratio) for 16-20 hours. Flow cytometry analysis of GFP mean fluorescence intensity revealed a 70% reduction in viability of HHT-treated THP-1 cells compared to a 25-28% decrease in DMSO-treated controls, demonstrating enhanced T cell-mediated killing of LILRB4-low tumor cells. (D) Co-culture of T cells with THP-1 cells pre-treated with 60 nM HHT or DMSO at a 1:1 ratio. The DMSO-treated group displayed a significant decrease in T cells. In contrast, the 60 nM HHT-treated group showed an elevated proportion of cytotoxic T cells, indicating that HHT mitigates the inhibitory effect of THP-1 cells on T cell proliferation. (E) Schematic representation of the LILRB4-mediated immune evasion mechanism in AML cells. The diagram depicts LILRB4 binding to its ligand APOE, initiating downstream SHP-2 phosphorylation and resulting in the release of ARG1. ARG1 then interacts with T cells, mediating immunosuppressive effects. This schematic was created using BioRender. (F) Western blot analysis of THP-1 cells treated with increasing concentrations of HHT. The results demonstrate that total SHP2 levels remain unchanged, while phosphorylated SHP2 (Y580) levels are reduced in a dose-dependent manner. (G-I) Western blot analysis of THP-1, NOMO-1, and MV4-11 cells treated with increasing concentrations of HHT. The results reveal a significant dose-dependent downregulation of ARG1, a key downstream effector protein in the LILRB4-mediated immune evasion pathway. (J) Western blot and flow cytometry analysis validating the efficiency of LILRB4 knockout in THP-1 cells. (K-L) THP-1 cells, with or without LILRB4 expression, were treated with 60 nM HHT or DMSO for 48 hours, washed, and re-cultured for an additional 48 hours. Conditioned medium was then collected for T cell culture. Results demonstrated that conditioned medium from DMSO-treated LILRB4^WT^ THP-1 cells significantly suppressed T cell proliferation compared to HHT-treated cells. In contrast, conditioned medium from LILRB4KO THP-1 cells had no impact on T cell proliferation, regardless of HHT or DMSO treatment.

To further elucidate whether the enhanced T cell-mediated cytotoxicity against monocytic acute myeloid leukemia (AML) cells by HHT is mediated through the HHT-induced reduction of LILRB4, and its subsequent impact on T cell proliferation, we conducted additional experiments. THP-1 cells were pre-treated with either HHT or DMSO. After treatment, the cells were thoroughly washed to remove any residual HHT and subsequently co-cultured with T cells (Fig. 6A). T cell proliferation was then assessed using flow cytometry. The results indicated that the population of T cells was significantly downregulated in co-cultures with DMSO-treated THP-1 cells compared to T cells cultured alone. This suggests that THP-1 cells, when not treated with HHT (and thus characterized by high levels of LILRB4), impair T cell proliferation. In contrast, T cells co-cultured with HHT-pretreated THP-1 cells (which exhibit low levels of LILRB4) demonstrated minimal changes in the proportions of T cell proliferation compared to T cells cultured alone (Fig. 6D and Fig. S6A). These findings indicate that HHT treatment, by reducing LILRB4 levels, diminishes the immunosuppressive capacity of THP-1 cells on T cell activity, thereby enhancing T cell-mediated cytotoxicity against monocytic AML cells.

Previous studies have demonstrated that LILRB4 suppresses T cell activity in AML cells through a signaling pathway involving APOE, LILRB4, SHP-2, uPAR, and arginase-1 (ARG1)^12^. Specifically, LILRB4 binds to the APOE ligand, activating downstream SHP-2 phosphorylation and ultimately leading to the release of ARG1, which binds to T cells and exerts immunosuppressive effects (Fig. 6E)^12^. To determine whether HHT impairs the immunosuppressive effects of THP-1 cells on T cells by reducing LILRB4 and its downstream signaling, we examined the changes in phosphorylated SHP-2 (p-SHP2) and ARG1 following HHT treatment. Western blot analysis revealed that HHT downregulated p-SHP2 (Y580) while total SHP-2 levels remained unchanged (Fig. 6F). Furthermore, treatment with HHT resulted in a significant decrease in ARG1 secretion in monocytic AML cell lines, including THP-1, NOMO-1, and MV4-11 cells (Fig. 6G-I). These findings indicate that HHT diminishes LILRB4 expression and its associated downstream signaling, including p-SHP2 and ARG1, thereby reducing the immunosuppressive effects of THP-1 cells on T cells.

To further determine whether HHT reduces the immunosuppressive effects of THP-1 cells on T cells through the reduction of LILRB4-mediated ARG1 release, we used CRISPR technology to knock out endogenous LILRB4 in THP-1 cells (Fig. 6J). THP-1 cells, both with and without LILRB4 expression, were treated with HHT or DMSO for 48 hours, then washed to eliminate residual HHT and re-cultured in fresh medium for an additional 48 hours before collecting the conditioned medium for T cell culture. The results showed that conditioned medium from DMSO-treated wild-type THP-1 cells (LILRB4^WT^) significantly inhibited T cell proliferation compared to medium from HHT-pretreated cells, indicating that HHT reduces the immunosuppressive effects of AML cells on T cells by diminishing ARG1 release in a LILRB4-dependent manner (Fig. 6K-L). In contrast, conditioned medium from LILRB4 knockout THP-1 cells (LILRB4^KO^) had no effect on T cell proliferation, regardless of HHT or DMSO pretreatment (Fig. 6K-L). These findings confirm that HHT primarily inhibits AML immune evasion by attenuating ARG1 release, a process highly dependent on LILRB4.

### HHT Suppress LILRB4-Mediated Immune Evasion in Acute Monocytic Leukemia

To examine whether HHT can suppress LILRB4-mediated immune evasion in vivo, we utilized humanized mouse xenograft models. Immune-deficient NCG mice were humanized by injecting with human peripheral blood mononuclear cells (PBMCs) isolated from healthy volunteers. Subsequently, either LILRB4 wild-type (LILRB4^WT^, WT) or knockout (LILRB4^KO^, B4KO) THP-1 cells were subcutaneously transplanted into the humanized mice, as illustrated in the experimental flowchart (Fig. 7A). Tumor formation was monitored following successful engraftment. As shown in Fig. 7B, tumors were significantly larger in mice transplanted with LILRB4^WT^ THP-1 cells compared to those transplanted with LILRB4^KO^ cells (Fig. 7B and Fig. S7A-C). Notably, HHT treatment significantly reduced tumor formation in mice bearing LILRB4^WT^ THP-1 cells, while it had no significant effect on tumor development in mice bearing LILRB4^KO^ cells (Fig. 7B and Fig. S7A-C). These results suggest that the inhibitory effect of HHT on tumor development is dependent on LILRB4.

**Figure 7.**
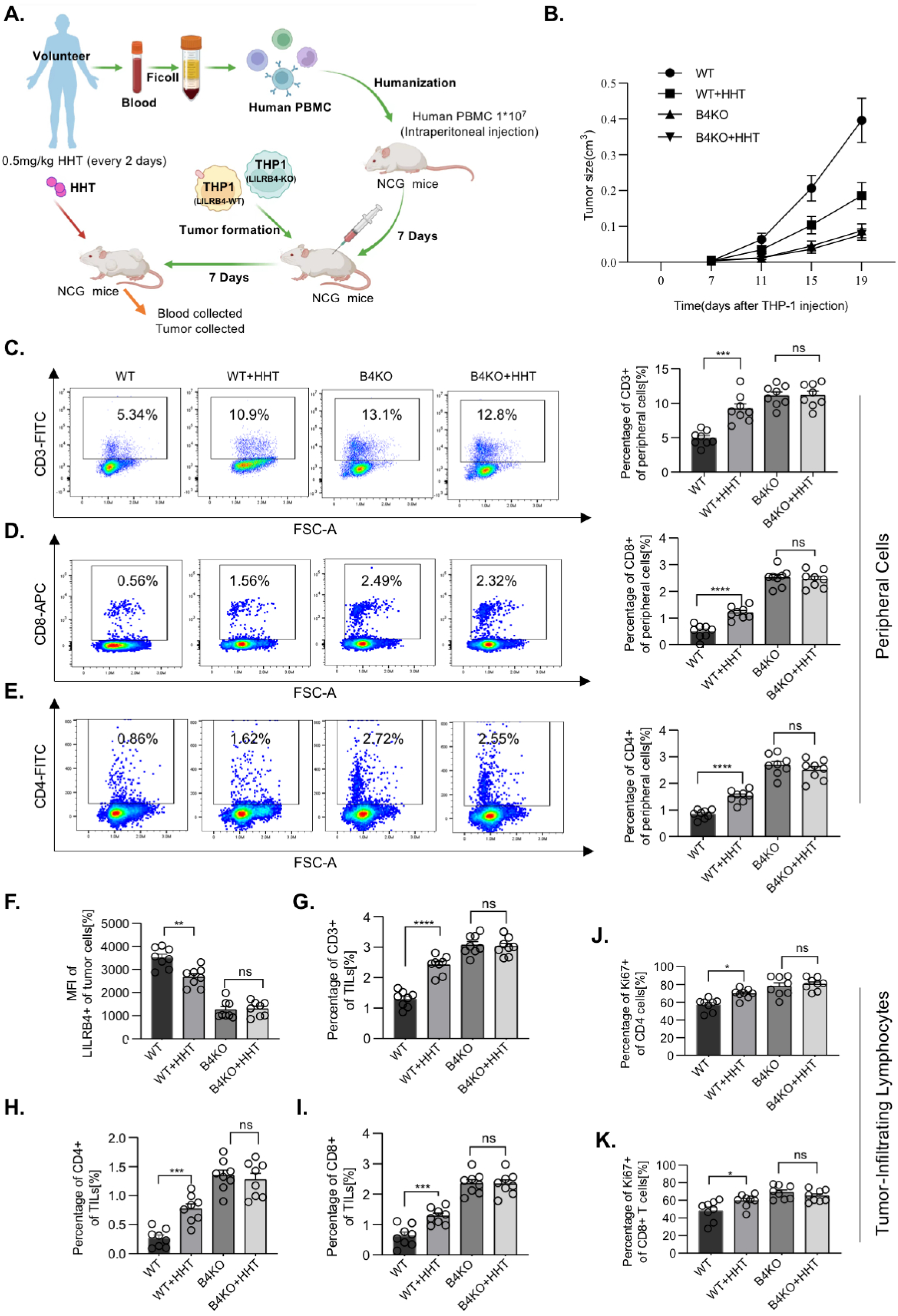
HHT Suppresses AML Tumor Growth in Humanized Mice. (A) Schematic representation of the experimental procedure: Human peripheral blood mononuclear cells (PBMCs) were isolated from volunteer blood using density gradient centrifugation by Ficoll. 1×10^7^ cells/per mice were then humanized in NCG mice via intraperitoneal injection. After seven days, tumor formation was induced by subcutaneous injection of THP-1 cells into NCG mice. 7 days later, mice were treated with either HHT (0.5 mg/kg every 2 days) or vehicle (PBS). Tumor growth was monitored for an additional 12 days. (B) The tumor size was calcultated (V=3/4*Πr^3^). (C-E) The graphs representing the percentages of CD3+ cells (C) and CD8+ cells (D) and CD4+ cell (E) among WT, WT+HHT, B4KO, and B4KO+HHT groups within the peripheral cells. WT: Mice subjected to subcutaneous injection of THP-1 cells expressing wild-type LILRB4 (LILRB4^WT^). WT+HHT: Mice subjected to subcutaneous injection of THP-1 cells expressing wild-type LILRB4 (LILRB4^WT^) and treated with the drug HHT. B4KO: Mice subjected to subcutaneous injection of THP-1 cells expressing knockout LILRB4 (LILRB4^KO^). B4KO+HHT: Mice subjected to subcutaneous injection of THP-1 cells expressing knockout LILRB4 (LILRB4^KO^) and treated with the drug HHT. (F) Bar graphs representing the expression of LILRB4 in tumor cells. (G-K) The graphs representing the percentages of CD3+ cells (G) and CD4+ cells (H) and CD8+ cell (I) and ki67+ of CD4+ cells (J) and ki67+ of CD8+ cells among WT, WT+HHT, B4KO, and B4KO+HHT groups within Tumor-infiltrating lymphocytes.

We further investigated the effects of HHT on T cell activity within the context of tumor development. Specifically, we quantified the percentage of human CD3^+^ T cells in the peripheral blood of mice from various experimental groups, including humanized mice transplanted with either LILRB4^WT^ or LILRB4^KO^ THP-1 cells, which were treated with HHT or PBS as a control. Given LILRB4’s established role in suppressing T cell activity to facilitate immune evasion by monocytic acute myeloid leukemia (AML), we hypothesized that HHT might counteract this immune suppression by decreasing LILRB4 levels. Consistent with our hypothesis, the percentage of human CD3^+^ T cells in peripheral blood was significantly higher in the LILRB4^KO^ group compared to the LILRB4^WT^ group (Fig. 7C and Fig. S7E). This finding underscores LILRB4’s critical role in AML-associated immune evasion via T cell suppression. Importantly, HHT treatment significantly increased the percentage of human CD3^+^ T cells in the peripheral blood of humanized mice transplanted with LILRB4^WT^ THP-1 cells, while having a minimal effect on the LILRB4^KO^ group (Fig. 7C and Fig. S7E). Similar trends were observed for the percentages of human CD8^+^ and CD4^+^ T cells under HHT treatment (Fig. 7D-E and Fig. S7F-G). Collectively, these data indicate that HHT treatment promotes T cell proliferation in a manner comparable to that seen in the LILRB4^KO^ group (Fig. 7C-E). Additionally, we observed that LILRB4 levels in humanized mice transplanted with LILRB4^WT^ THP-1 cells decreased during HHT treatment (Fig. 7F and Fig. S7D), consistent with our previous findings at the cellular level.

As tumor-infiltrating lymphocytes (TILs) are essential for tumor suppression, we examined the status of T cells within the tumor. An increase in the percentage of infiltrating T cells (CD3^+^ T cells) in the tumor was observed in the LILRB4^KO^ group. Moreover, HHT treatment significantly increased the percentage of human CD3^+^ T cells in TILs of humanized mice transplanted with LILRB4^WT^ THP-1 cells, while having a minimal effect on the LILRB4^KO^ group, similar to the observations in peripheral blood (Fig. 7G and Fig. S7E). Similarly, the percentages of CD4^+^ and CD8^+^ T cells in TILs were increased under HHT treatment (Fig. 7H-I and Fig. S7F-G). These tumor-infiltrating T cells exhibited enhanced proliferative capacities, as evidenced by increased Ki67 expression (Fig. 7J-K and Fig. S7H-I). These findings suggest that LILRB4 knockdown or downregulation by HHT treatment may have an anti-tumor effect. Overall, these results together support the notion that HHT inhibits tumor formation by suppressing immune evasion in AML, likely through the reduction of LILRB4 levels. This mechanistic insight highlights HHT’s potential as a therapeutic agent for AML by enhancing immune surveillance and overcoming immune evasion mediated by LILRB4.

To validate the clinical relevance of our findings, we isolated monocytic AML cells from patients with subtypes M4 and M5 and cultured them in RPMI 1640 medium. These cells were treated with varying concentrations of HHT (0–160 nM) for 2 days, and the expression levels of LILRB4, FTO, and MLL were analyzed using Western blot. The results showed that HHT significantly reduced the expression of these proteins, consistent with our findings in THP-1 cells (Fig. 8A-D). This suggests that HHT can effectively lower LILRB4 levels in AML patients with subtypes M4 and M5. We next conducted a T cell co-culture assay to assess HHT’s impact on the immunosuppressive effects of monocytic AML cells from AML-M4/5 patients. The monocytic AML cells from patients were pre-treated with HHT or DMSO for 48 hours, washed, and re-cultured in fresh medium for another 48 hours. The conditioned medium from these treated cells was then collected and used to culture T cells. The results indicated that the conditioned medium from HHT-treated AML cells significantly reduced the suppression of T cell activity compared to the DMSO control (Fig. 8E). These findings support the potential clinical application of HHT’s anti-immunosuppressive effects and confirm its efficacy against LILRB4 in AML patients.

**Figure 8.**
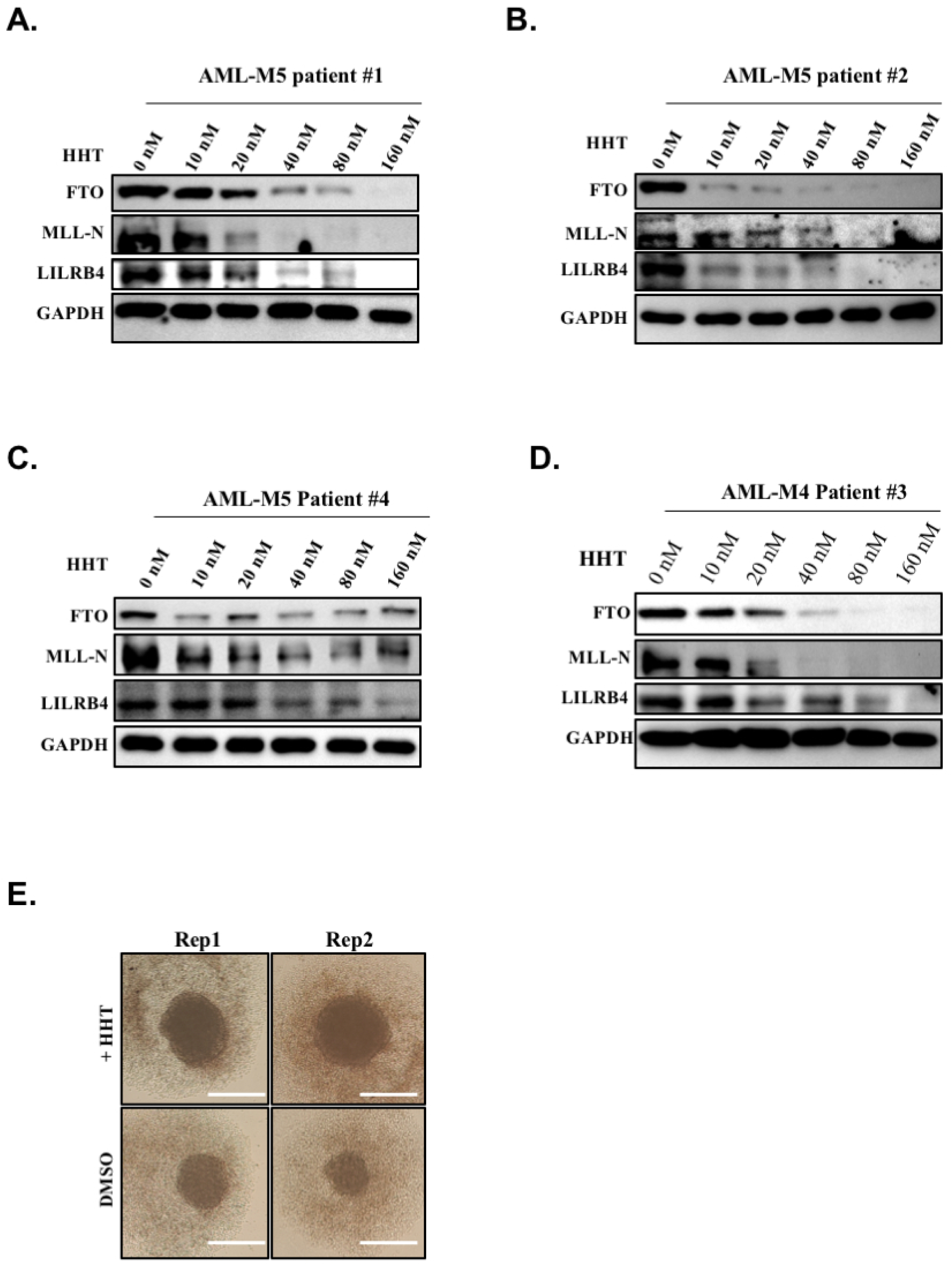
HHT Suppresses LILRB4 Expression in Primary Cells from AML-M4/5 Clinical Samples. (A-D) Western blot analysis of primary cells from an AML-M4/M5 patient treated with a range of 0-160 nM HHT reveals a dose-dependent decrease in the expression of LILRB4, FTO, and MLL proteins. (A) Primary cells from an AML-M5 patient were pre-treated with HHT for 48 hours, washed to remove the residual HHT, and then re-cultured in fresh medium for another 48 hours. T cells were cultured with conditioned media collected from the above primary cells. The results demonstrate that HHT treatment significantly enhances T cell proliferation, as evidenced by the increased formation of T cell colonies.

## Discussion

In this study, we investigated the role of HHT in targeting the immune evasion mechanisms utilized by acute monocytic leukemia (AML). Our findings reveal that HHT promotes the degradation of FTO, which subsequently leads to decreased expression of LILRB4, an immune inhibitory receptor specifically expressed on monocytic AML cells that facilitates immune escape. This discovery provides insights into the potential mechanisms through which HHT may enhance the efficacy of immune responses against AML.

Acute monocytic leukemia (AML-M5) is an aggressive malignancy characterized by a poor prognosis, primarily due to its capacity to evade immune surveillance. Our study elucidates that HHT can effectively counteract this immune evasion by targeting the FTO/m6A/LILRB4 signaling pathway. We demonstrate that HHT significantly downregulates LILRB4 expression at both the RNA and protein levels, suggesting that its effects extend beyond its established role as a protein synthesis inhibitor. Mechanistically, HHT treatment induces a marked reduction in FTO protein levels through a GSK3β-dependent degradation pathway mediated by the ubiquitin-proteasome system. FTO, an RNA demethylase, is pivotal in modulating gene expression by influencing RNA stability and translation efficiency. By degrading FTO, HHT alters the balance of m6A methylation, leading to the downregulation of key downstream targets, including MLL1 and LILRB4. This mechanism highlights the intricate interplay between RNA modifications and immune evasion in leukemia, underscoring the therapeutic potential of HHT for managing AML. Our findings provide novel insights into how HHT can enhance immune responses by targeting the FTO/m6A/LILRB4 pathway, offering a promising strategy for overcoming immune evasion in AML.

The downregulation of LILRB4 following HHT treatment highlights the potential of targeting this pathway to enhance immune-mediated clearance in acute myeloid leukemia (AML). LILRB4 is implicated in immunosuppressive signaling that facilitates immune evasion by leukemia cells. Our comprehensive studies, encompassing in vitro assays, mouse xenograft models, and clinical samples from AML-M5 patients, demonstrate that HHT reduces LILRB4 expression and increases the susceptibility of AML-M5 cells to T cell-mediated cytotoxicity. These findings suggest that HHT may reinvigorate the anti-tumor immune response in AML-M5 patients. The observed increase in immune activation markers further supports the potential of HHT to enhance immunotherapeutic efficacy. This discovery emphasizes the therapeutic potential of HHT in AML immunotherapy.

By reducing LILRB4 levels, HHT can restore the ability of immunocompetent cells, such as T cells, to recognize and eliminate leukemic cells. This is particularly significant given the recognized role of the immune microenvironment in acute myeloid leukemia (AML) progression and therapy resistance. For instance, Decitabine (DAC), an epigenetic drug used to treat myelodysplastic syndromes (MDS) and AML, has been reported to increase LILRB4 expression by promoting FTO expression, leading to acquired resistance during long-term treatment of human AML cells^44,58^. In this study, we demonstrate that HHT reduces LILRB4 levels by promoting FTO degradation. This finding provides a rationale for combining HHT and DAC to treat AML patients, potentially overcoming DAC-induced resistance while suppressing AML progression. This discovery emphasizes the therapeutic potential of HHT not only as a chemotherapeutic agent but also as an immunomodulatory agent capable of reinvigorating immune responses against AML, and may even be effective against those AML cases that are immunologically refractory.

While our study has provided valuable insights into the mechanisms by which HHT promotes FTO degradation to suppress LILRB4-mediated immune evasion in acute monocytic leukemia (AML-M5), several questions remain unresolved and necessitate further investigation. Although we have identified MLL1 and its associated H3K4me3 histone modification as intermediaries in FTO’s regulation of LILRB4, we have not thoroughly examined how FTO regulates MLL1 levels in greater detail in this study. Existing literature suggests that FTO can modulate MLL1 levels via the ASB2 protein, a ubiquitin ligase that facilitates the degradation of MLL1^54,59^. Therefore, it is essential to determine whether HHT’s regulation of MLL1 through FTO is dependent on this mechanism. Additionally, our research indicates that HHT treatment reduces GSK-3β phosphorylation levels. Prior studies have shown that HHT targets the PI3K/AKT/GSK3β/Slug signaling pathway to suppress the progression of hepatocellular carcinoma^60^. However, it remains unclear which kinases are the primary targets of HHT in vivo and how HHT directly targets these kinases to regulate this process in AML. Further research is needed to identify the primary kinases targeted by HHT in vivo and to understand how HHT directly interacts with these kinases to modulate the PI3K/AKT/GSK3β/Slug signaling pathway in AML. These investigations will provide a more comprehensive understanding of HHT’s therapeutic potential and may lead to the development of more effective treatment strategies for acute monocytic leukemia.

In conclusion, our study demonstrates that HHT is a promising agent for mitigating immune evasion in acute monocytic leukemia by promoting FTO degradation and reducing LILRB4 expression. While our findings highlight the therapeutic potential of HHT and the importance of targeting immune escape mechanisms in aggressive leukemias, we acknowledge that HHT’s antitumor effects likely extend beyond its action on the FTO/m6A/LILRB4 pathway. We propose that inhibiting immune evasion via this pathway is a key mechanism underlying HHT’s antitumor activity. Our research opens new avenues for therapeutic strategies that integrate epigenetic regulation and immune response to improve AML treatment efficacy. Further clinical studies are needed to validate these findings, evaluate HHT’s role in enhancing CD8^+^ T cell-based immunotherapy for AML, and explore its full potential in combination therapies.

## Acknowledgements

This work was supported by the National Natural Science Foundation of China (82000171 to F.H., 32270638 and 32100464 to S.C.), the National Key Research and Development Program of China (2023YFE0118000 to S.C.), the Shenzhen Science and Technology Innovation Commission (JCYJ20230807091204009 to S.C.), Fujian Provincial Natural Science Foundation of China (2020J05291 to F.H. and 2022J01051 to S.C.), the Fundamental Research Funds for the Central Universities (20720220122 to S.C.), the Nanqiang Outstanding Young Talents Program from Xiamen University (to S.C.), and Joint Project of Pinnacle Disciplinary Group, the Second Affiliated Hospital of Chongqing Medical University (2024304 to Q.L.), The graphical abstract was generated with BioRender. We sincerely thank Dr. Zunling Li for providing essential materials and guidance.

## Ethics approval and consent to participate

The animal study was reviewed and approved by the Laboratory Animal Welfare and Ethics Committee of Xiamen University. All patients provided written informed consent. The experiments were approved by the Medical Ethics Committee of Xiamen University (No: 82000171) and conducted in accordance with national standards for Institutional Animal Care and Use.

## Author contributions

S.C. and F.H. conceived the project. F.H., M.Z., X.L., and L.J. designed and performed most of the experiments with help from W.S., P.C., X.H., C.X., M.J., D.H., B.Z., S.H., and C.Y.; J.W. helped perform the experiments shown in Figure 7, Supplementary Figure 7, and analyzed the data; J.Z., R.G., and Q.L. provided critical materials or instructions; S.C. and F.H. wrote the manuscript; S.C. supervised the project with J.W. and Q.L.

