## supplemental Files for "Homoharringtonine Promotes FTO Degradation to Suppress LILRB4-Mediated Immune Evasion in Acute Monocytic Leukemia"

**This PDF file includes:**

**Figs. S1 to S7**

### STAR★METHODS

#### KEY RESOURCES TABLE

| REAGENT or RESOURCE | SOURCE | IDENTIFIER |
|---|---|---|
| **Antibodies** | | |
| Flag Mouse mAb | Sigma-Aldrich | Cat No. F1804 |
| H3K27me3 Rabbit mAb | Cell Signaling Technology | Cat No. 9733S |
| Histone H3 Rabbit mAb | Cell Signaling Technology | Cat No. 4499S |
| EZH2 Rabbit mAb | Cell Signaling Technology | Cat No. 5246S |
| SUZ12 Rabbit mAb | Cell Signaling Technology | Cat No. 3737S |
| EED Rabbit mAb | Cell Signaling Technology | Cat No. 85322S |
| SHP-2 Rabbit mAb | Cell Signaling Technology | Cat No. 3397S |
| Phospho-SHP-2 (Tyr580) Antibody | Cell Signaling Technology | Cat No. 3703S |
| MLL1-N Rabbit mAb | Cell Signaling Technology | Cat No. 14689S |
| Phospho-GSK-3β (Ser9) Rabbit mAb | Cell Signaling Technology | Cat No. 9323S |
| Anti-mouse IgG, HRP-linked Antibody | Cell Signaling Technology | Cat No. 7076S |
| Anti-rabbit IgG, HRP-linked Antibody | Cell Signaling Technology | Cat No. 7074S |
| GAPDH Monoclonal Antibody | Invitrogen | Cat No. MA1-16757 |
| DOT1L antibody (pAb) | Active Motif | Cat No. 39954 |
| LILRB4 Rabbit mAb | R&D | Cat No. MAB24251-SP |
| PSIP1 Polyclonal antibody | Proteintech | Cat No. 25504-1-AP |
| METTL14 Polyclonal antibody | Proteintech | Cat No. 26158-1-AP |
| METTL3 Polyclonal antibody | Proteintech | Cat No. 15073-1-AP |
| IGF2BP2 Polyclonal antibody | Proteintech | Cat No. 11601-1-AP |
| m6A Monoclonal antibody | Proteintech | Cat No. 68055-1-Ig |
| USP7 Monoclonal antibody | Proteintech | Cat No. 66514-1-Ig |
| FTO Polyclonal antibody | Proteintech | Cat No. 27226-1-AP |
| RARA Polyclonal antibody | Proteintech | Cat No. 10331-1-AP |

| | | |
|---|---|---|
| c-MYC Polyclonal antibody | Proteintech | Cat No. 10828-1-AP |
| ubiquitin Polyclonal antibody | Proteintech | Cat No. 10201-2-AP |
| β-Actin Recombinant antibody | Proteintech | Cat No. 81115-1-RR |
| GSK3β Rabbit pAb | Abclonal | Cat No. A2081 |
| H3K79me2 Rabbit mAb | Abclonal | Cat No. A22086 |
| H3K4 me3 Rabbit mAb | Abclonal | Cat No. A22146 |
| MEN1 Rabbit mAb | Abclonal | Cat No. A3395 |
| WDR5 Rabbit mAb | Abclonal | Cat No. A3259 |
| ASH2L Rabbit mAb | Abclonal | Cat No. A4892 |
| RBBP5 Rabbit mAb | Abclonal | Cat No. A23124 |
| KMT2A Rabbit pAb | Abclonal | Cat No. A1435 |
| FITC Mouse anti-Human CD3 mAb | Abclonal | Cat No. A22794 |
| ABflo® 594 Rabbit anti-Human CD4 mAb | Abclonal | Cat No. A25282 |
| ABflo® 647 Rabbit anti-Human/Monkey CD8a mAb | Abclonal | Cat No. A23346 |
| ARG1 Rabbit Monoclonal Antibody | HUABIO | Cat No. ET1605-8 |
| **REAGENT or RESOURCE** | | |
| MinElute PCR Purification Kit | QIAGEN | Cat. No. 28004 |
| AG RNAex Pro Reagent | Accurate Biotechnology | Cat No. AG21101 |
| Evo M-MLV Reverse Transcriptase Kit | Accurate Biotechnology | Cat No. AG11728 |
| SYBR Green Pro Taq HS qPCR Kit | Accurate Biotechnology | Cat No. AG11739 |
| PBMC Isolation Kit | Solarbio | Cat No. P8610 |
| Human CD3/CD28 T cell Activation Beads Kit | Proteintech | Cat No. KMS310 |
| Active Recombinant Human IL-2 Protein | Abclonal | Cat No. RP01039 |
| Lenti-X concentrator | Takara | Cat No. 631231 |
| Fetal bovine serum (FBS) | Vivacell | Cat No. 2224109 |
| Anti-DYKDDDDK G1 Affinity Resin | GenScript | Cat. No. L00432 |

| | | |
|---|---|---|
| High Affinity Ni-Charged Resin | GenScript | Cat. No. L00666 |
| **Chemicals, Peptides, and Recombinant Proteins** | | |
| Homoharringtonine (HHT) | APExBIO | Cat. No. N1504 |
| FB23-2 | MedChemExpress | Cat. No. HY-127103 |
| SB 415286 | TargetMol | Cat. No. 264218-23-7 |
| a-Ketoglutaric acid | Macklin | Cat. No. K81223 |
| L-Ascorbic acid | Sangon Biotech | Cat. No. A610021 |
| Ammonium iron (II) sulfate hexahydrate | Sangon Biotech | Cat. No. A600065 |
| Puromycin | Yeasen Biotechnology | Cat No. 60210ES25 |
| G418 Sulfate | Yeasen Biotechnology | Cat. No. 60220ES03 |
| Blasticidin | Yeasen Biotechnology | Cat. No. 60218ES10 |
| Penicillin Streptomycin | Pricella | Cat No. PB180120 |
| Amersham Hybond-N+ membrane | Cytiva | Cat. # RPN303B |
| cOmplete™, EDTA-free Protease Inhibitor Cocktail | Roche | Cat. # 04693132001 |
| PVDF membranes | Bio-Rad | Cat No. 1620177 |
| PEI | Yeasen Biotechnology | Cat No. 40816ES03 |
| Formaldehyde | Sigma | Cat No. F1635 |
| IPTG | INALCO | Cat No. 1758-1400 |
| DTT | INALCO | Cat No. 1758-9030 |
| **Experimental Models: Cell Lines** | | |
| Human cell line: HEK293 | ATCC | CRL-1573 |
| Human cell line: THP-1 | ATCC | TIB-202 |
| Human cell line: MV4-11 | ATCC | CRL-9591™ |
| Human cell line: NOMO-1 | DSMZ | ACC 542 |
| THP-1: Sg0 | This Paper | N/A |
| THP-1: LILRB4-KO | This Paper | N/A |
| THP-1: MLL1-KO | This Paper | N/A |
| THP-1: ShNC | This Paper | N/A |

| | | |
|---|---|---|
| THP-1: ShFTO-1 | This Paper | N/A |
| THP-1: ShFTO-2 | This Paper | N/A |
| MV4-11: ShNC | This Paper | N/A |
| MV4-11: ShFTO-1 | This Paper | N/A |
| MV4-11: ShFTO-2 | This Paper | N/A |
| THP-1: ShGSK3β | This Paper | N/A |
| THP-1: MLL1-KO + FTO$^{WT}$ | This Paper | N/A |
| THP-1: MLL1-KO + FTO$^{Mut}$ | This Paper | N/A |
| THP-1: ShMLL1-1 | This Paper | N/A |
| THP-1: ShMLL1-2 | This Paper | N/A |
| **Recombinant DNA** | | |
| pCDH-3×Flag-FTO | This Paper | N/A |
| pCDH-3×Flag-FTO $^{H231A/D233A}$ | This Paper | N/A |
| pLKO.1-shFTO-1 | This Paper | N/A |
| pLKO.1-shFTO-2 | This Paper | N/A |
| pLKO.1-shMLL1-1 | This Paper | N/A |
| pLKO.1-shMLL1-2 | This Paper | N/A |
| pLKO.1-shNC | This Paper | N/A |
| pLKO.1-shGSK3β | This Paper | N/A |
| LentiCRISPR v2 | This Paper | Addgene #52961 |
| psPAX2 | This Paper | Addgene #12260 |
| VsVg | This Paper | Addgene #8454 |
| pCDNA3.0-Myc-GSK3β | This Paper | N/A |
| pCDNA3.0-FLAG-FTO | This Paper | N/A |
| pCDNA3.0-HA-Ub | This Paper | N/A |

#### RESOURCE AVAILABILITY

#### EXPERIMENTAL MODEL AND SUBJECT DETAILS

##### Cell Cultures

THP-1, MV4-11, and NOMO-1 cells were cultured in RPMI 1640 medium supplemented with 10% fetal bovine serum (FBS) (Vivacell, 2224109) and 1% penicillin-streptomycin (Pricella, PB180120). HEK293T cells were cultured in Dulbecco's Modified Eagle Medium (DMEM) supplemented with 10% FBS (Vivacell, 2224109) and 1% penicillin-streptomycin (Pricella, PB180120). All cells were maintained in a humidified incubator at 37°C with 5% $CO_2$.

##### Clinical Specimens

Clinical peripheral blood samples were obtained from patients diagnosed with M4/M5 acute myeloid leukemia (AML) at Zhongshan Hospital Affiliated to Xiamen University. The study was approved by the Institutional Ethics Committee of Zhongshan Hospital of Xiamen University, and informed consent was obtained from all participants. Mononuclear cells were isolated from the peripheral blood samples using density gradient centrifugation with lymphocyte separation medium. The isolated cells were treated with PBS and red blood cell lysis solution to obtain acute monocytic leukemia cells. These cells were then cultured in RPMI-1640 medium supplemented with 10% fetal bovine serum (FBS) at a density of $5 \times 10^5$ cells/mL in six-well plates. The cultured cells were treated with Homoharringtonine (HHT) at concentrations ranging from 0 to 160 nM for 48 hours. Following treatment, the cells were collected and lysed for Western blot analysis to examine the relevant indicators. All research was conducted in accordance with government policies and the Helsinki Declaration. Demographic information, including the gender and age of the patients, is provided in Table S3.

**Animal Studies**

Male NCG mice (4-5 weeks old) were obtained from GemPharmatech Co., Ltd, China, and housed in the animal laboratory of Xiamen University under a 12-hour light/12-hour dark cycle. Initially, lymphocytes were isolated from peripheral blood samples collected from healthy volunteers and intravenously injected into the mice. After seven days, the proportion of T cells among lymphocytes was assessed, with a threshold of over 80% indicating successful model establishment. Subsequently, human THP-1 ($LILRB4^{WT}$) and THP-1 ($LILRB4^{KO}$) cells were subcutaneously injected into the mice. Once tumors reached the size of a soybean, mice were treated with either PBS or 0.5 mg/kg Homoharringtonine (HHT) until the tumor volume approached 1 $cm^3$. The mice were then euthanized, and blood, tumor, and spleen tissues were collected. Flow cytometry was employed to determine whether the inhibitory effect of HHT on acute myeloid leukemia (AML) cells in suppressing tumor immune escape was dependent on the presence of LILRB4. All animal experiments were approved by the Xiamen Animal Care and Use Committee and conducted in accordance with relevant guidelines and regulations.

#### METHOD DETAILS

**Expression and Purification of Human FTO.**

The cDNA sequence encoding human FTO (residues 32-505) was cloned into the pET-28a vector, incorporating an N-terminal His6 tag. The recombinant plasmid was transformed into *E. coli* Rosetta 2 (DE3) cells. For protein expression, transformed cells were cultured in Luria-Bertani (LB) medium at 37°C until the optical density at 600 nm (OD600) reached 0.6. Protein expression was then induced by adding 0.5 mM isopropyl-1-thio-β-D-galactopyranoside (IPTG) and incubating the culture at 20°C for 18 hours. After induction, the cells were harvested by centrifugation and resuspended in lysis buffer containing 50 mM Tris-HCl (pH 8.0), 500 mM NaCl, 5% glycerol, 2 mM β-mercaptoethanol, and 1 mM phenylmethylsulfonyl fluoride (PMSF). The cell suspension was lysed by sonication, and the lysate was cleared by centrifugation at 4°C. The supernatant was applied to a Ni-NTA resin column equilibrated with lysis buffer. The resin was washed extensively with Ni-NTA wash buffer (50 mM Tris-HCl, pH 8.0, 500 mM NaCl, 20 mM imidazole, 5% glycerol, and 2 mM β-mercaptoethanol). The His6-tagged FTO protein was eluted with the wash buffer supplemented with 300 mM imidazole. Further purification was achieved using Mono Q ion exchange chromatography. Fractions containing the target protein were pooled and concentrated. The concentrated protein was further purified by size exclusion chromatography (SEC) using a Superdex 200 column equilibrated with SEC buffer (20 mM Tris-HCl, pH 8.0, 100 mM NaCl, and 2 mM DTT). The eluate was analyzed by SDS-PAGE to confirm the purity of the protein. The purified protein was then concentrated and stored at -80°C for subsequent experiments.

**m6A Dot-Blot Assay**

To evaluate the effect of Homoharringtonine (HHT) on m6A RNA methylation in THP-1 cells, we treated the cells with either DMSO or various concentrations of HHT for 48 hours. Total RNA was extracted using TRIzol reagent (Invitrogen) following the manufacturer's protocol. The RNA was then diluted in RNase-free water and denatured by heating at 95°C for 5 minutes. The denatured RNA samples were dotted onto an Amersham Hybond-N+ membrane (Cytiva, RPN303B) using a Bio-Dot Apparatus (Bio-Rad). The RNA samples were cross-linked to the membrane by exposure to 254 nm UV light for 5 minutes. The membrane was blocked with 5% non-fat milk in Tris-buffered saline with Tween-20 (TBST) at room temperature for 1 hour. Subsequently, the membrane was incubated overnight at 4°C with an anti-m6A monoclonal antibody (Proteintech, Cat No. 68055-1-Ig, diluted 1:1000). After washing with TBST, the membrane was incubated with horseradish peroxidase (HRP)-conjugated horse anti-mouse IgG secondary antibody (Cell Signaling Technology, #7076) for 1 hour at room temperature. The m6A-modified RNA was detected using the ECL Prime Western Blot Detection Reagent (Bio-Rad) according to the manufacturer's instructions.

**Immunoblotting (Western Blot)**

THP-1, MV4-11, and NOMO-1 cells were treated with HHT for 48 hours. Post-treatment, cells were lysed in RIPA Lysis Buffer at 4°C. Protein concentrations were quantified using a BCA Protein Assay Kit and normalized to ensure equal loading. Equivalent amounts of whole-cell lysates were subjected to SDS-PAGE and transferred onto PVDF membranes (Bio-Rad, Cat# 1620177). Membranes were blocked with 5% non-fat milk in Tris-buffered saline with Tween-20 (TBST) at room temperature for 1 hour. Subsequently, membranes were incubated overnight at 4°C with primary antibodies specific to the proteins of interest. After washing with TBST, membranes were incubated for 1 hour at room temperature with secondary antibodies: HRP-conjugated horse anti-mouse IgG (Cell Signaling Technology, Cat# 7076) or HRP-conjugated goat anti-rabbit IgG (Cell Signaling Technology, Cat# 7074). Detection was carried out using Clarity Western ECL Substrate (Bio-Rad, Cat# 170-5061) according to the manufacturer's instructions. Chemiluminescent signals were captured using a ChemiDoc Imaging System (Bio-Rad).

**Detection of FTO Enzymatic Activity by PAGE-Based Assays**

To assess the effect of Homoharringtonine (HHT) on the enzymatic activity of FTO in vitro, we conducted a PAGE-based assay. The assay utilized a methylated 49-nucleotide single-stranded DNA (ssDNA) substrate containing a DpnII cleavage site (5′-TAGACATTGCCATTCTCGATAGG(dm6A)TCCGGTCAAACCTAGACGAATTCCA-3′). The reaction mixtures consisted of 50 mM Tris-HCl (pH 7.5), 1 μM ssDNA, 1 μM truncated FTO (FTO△N31), 300 μM α-ketoglutaric acid, 280 μM $(NH4)_2Fe(SO4)_2$, 2 mM L-ascorbic acid, and varying concentrations of HHT. The reactions were incubated at room temperature for 2 hours and subsequently terminated by heating. The ssDNA was annealed to its complementary strand to facilitate DpnII digestion. The digestion products were resolved on a 15% non-reducing PAGE gel and visualized using Gel-Red staining. Band intensities were quantified to evaluate the enzymatic activity of FTO.

**Real-Time Quantitative PCR (RT-qPCR)**

Total RNA samples were isolated with the AG RNAex Pro reagent (Accurate Biology, AG21101) and subjected to reverse transcription using the Evo M-MLV Reverse transcriptase Kit (Accurate Biology, AG11728). RT-qPCR reactions were performed with SYBR Green Pro Taq HS qPCR Kit (Accurate Biology, 1G11739) with a 0.1 mL qPCR 96 well clear plate (Accurate Biology, AG12116) in an AriaMx Real-time PCR System (Agilent Technologies). Gene expression was calculated using the comparative ΔΔCT method with GAPDH expression used for normalization. Each reaction was run in triplicates. The primers used in this study are listed in Table S1.

**Isolation and Culture of Human Primary Lymphocytes**

Human peripheral blood was obtained from healthy volunteers and diluted 1:1 with phosphate-buffered saline (PBS, 1×). An equal volume of lymphocyte separation medium was added to a new centrifuge tube, and the diluted peripheral blood was carefully layered on top of the separation medium without mixing. The mixture was then centrifuged at 400-500 g for 30-40 minutes at room temperature. After centrifugation, the lymphocyte layer was carefully transferred to a new centrifuge tube, washed with 10 mL of PBS, and centrifuged at 300-400 g for 10 minutes. The supernatant was discarded, and the cells were washed twice with PBS by centrifugation at 300-400 g for 10 minutes each. Following centrifugation, residual red blood cells were removed using red blood cell lysis buffer and then centrifuged at 300-400 g for 5-10 minutes to eliminate the lysis buffer at room temperature. The resulting lymphocytes were washed once with 5 mL of PBS by centrifugation at 300-400 g for 10 minutes. The isolated lymphocytes were resuspended in RPMI-1640 complete medium supplemented with 10% fetal bovine serum (FBS), 1% penicillin-streptomycin, and interleukin-2 (IL-2), and cultured in a 12-well plate at a density of $1 \times 10^6$ cells/mL. The isolated lymphocytes can be stained with anti-CD3 antibody and purified by fluorescence-activated cell sorting (FACS) to obtain T cells.

**Co-culture Assay with AML Cells and T Cells**

To explore the interaction between acute myeloid leukemia (AML) cells and T cells, we performed a co-culture experiment. Peripheral blood mononuclear cells (PBMCs) were isolated from healthy donor peripheral blood using a lymphocyte separation solution kit (Solarbio, P8610). T lymphocytes were subsequently separated and collected from these PBMCs. The T cells were cultured in RPMI 1640 medium supplemented with a human CD3/CD28 T cell activation beads kit (Proteintech, KMS310) and 50 U/ml of active recombinant human IL-2 protein (Abclonal, RP01039). In parallel, THP-1 cells were infected with a lentivirus designed to constitutively express GFP. GFP-positive cells were selected and treated with either 30 nM Homoharringtonine (HHT) or DMSO for 48 hours. Post-treatment, the GFP-positive cells were harvested, washed once with fresh medium, and resuspended in fresh medium. These HHT-pretreated GFP-positive cells (20,000 cells/well) were then co-cultured with T cells (20,000 cells/well) in 24-well plates for 12-16 hours. Following the co-culture period, the cells were stained with anti-Human CD4 monoclonal antibody (mAb) (Abclonal, A25282) and Rabbit anti-Human/Monkey CD8a mAb (Abclonal, A23346). The number of GFP-positive cells and the ratios of CD4/CD8 differentiation were analyzed using flow cytometry with FlowJo V10 software.

**Lentivirus Production and Delivery**

Lentivirus were produced for the pCDH-EF1-3×FLAG-FTO WT or mutant plasmids, pLKO.1-shFTO-1, pLKO.1-shFTO-2, pLKO.1-MLL1-1, pLKO.1-MLL1-2, and the control pLKO.1-shNC. The packaging was performed using psPAX2 (Addgene, #12260) and VsVg (Addgene, #8454). Co-transfection into 293T cells was carried out with 2.25 μg of psPAX2, 0.75 μg of VsVg, and 3 μg of the respective constructs using PEI (Yeasen Biotechnology, 40816ES03). After 48 hours of transfection, the virus-containing medium was collected and concentrated using a Lenti-X concentrator (Takara, 631231). The viral pellet was collected by centrifugation and re-suspended in culture medium for subsequent infection of AML cells. To select positively infected cells, 1.5 mg/ml of puromycin (Yeasen Biotechnology, 60210ES25) was added to the cultures 48 hours post-infection. The oligonucleotides used in this study are listed in Table S2

**CRISPR-Cas9-Based Genome Editing**

The CRISPR/Cas9 system was employed to target MLL1 and LILRB4 in THP-1 and MV4-11 cells. Specific sgRNA sequences were designed using the online tool available at CRISPR Design Tool and cloned into the LentiCRISPR v2 vector (Addgene, #52961). Lentivirus carrying the desired sgRNA for MLL1 or LILRB4 was then used to establish stable cell lines. HEK293T cells were co-transfected with the packaging plasmid psPAX2 (Addgene, #12260), envelope plasmid VsVg (Addgene, #8454), and the relevant constructs for 48 hours. Following transfection, the virus-containing medium was harvested and concentrated using a Lenti-X concentrator (Takara, 631231). Stable cell lines were developed by selecting with puromycin, and monoclonal cells were further cultured. The infection efficiency was verified through western blot analysis and PCR amplification of genomic DNA, followed by sequencing. The sgRNA sequences used in this study are listed in Table S2.

**Flow Cytometry Analysis**

Flow cytometry analysis was performed to investigate the effects of homoharringtonine (HHT) on the expression of LILRB4 in THP-1 cells. THP-1 cells were treated with HHT or DMSO at specified concentrations for 48 or 72 hours. The antibodies used in this study included PE-conjugated anti-human CD85k (ILT3) Antibody (Biolegend, 333008) and anti-mouse IgG1, κ Isotype Control Antibody (Biolegend, 40014) as a control for non-specific binding. For surface staining, cells treated with various concentrations of HHT were harvested, washed twice with chilled phosphate-buffered saline (PBS), and then stained with PE-conjugated anti-human CD85k (ILT3) antibody or anti-mouse IgG1, κ Isotype Control Antibody. The cells were incubated for 15-20 minutes at 4°C in the dark before FACS analysis. For cell cycle analysis, cells treated with different concentrations of HHT were collected by centrifugation, fixed with 1 mL of 70% ice-cold ethanol, and stored at -20°C for at least 2 hours. After fixation, the cells were washed with PBS and stained using the Cell Cycle and Apoptosis Analysis Kit (Yeasen, 40301ES50). Each sample was resuspended in 500 µL of staining solution containing 10 µL propidium iodide (PI) and 10 µL RNase A, followed by incubation at 37°C for 30 minutes in the dark. The stained cells were then analyzed directly on a flow cytometer. Data were analyzed using FlowJo V10 software.

**Co-immunoprecipitation**

To explore the interaction between GSK-3β and FTO, we conducted a co-immunoprecipitation experiment. The full-length GSK-3β and FTO genes were cloned into the pCDNA3.0 vector, each tagged with an N-terminal MYC and FLAG tag, respectively. HEK293T cells were transiently transfected with these plasmids using standard protocols. After 48 hours, the transfected cells were harvested and lysed in IP lysis buffer (50 mM Tris, 150 mM NaCl, 2 mM EDTA, 0.5% NP-40, and 5 mM MgCl2; pH 7.4) supplemented with a protease inhibitor cocktail (Roche). The lysates were clarified by centrifugation at 14,000 × g for 15 minutes at 4°C. FTO and its associated proteins were captured using Anti-FLAG M2 beads (Sigma-Aldrich) and washed three times with a buffer containing 50 mM Tris-HCl (pH 8.0), 150 mM NaCl, 2 mM DTT, 5 mM MgCl2, and 0.1% NP-40. The isolated FTO and associated proteins were then analyzed by Western blot using specific antibodies.

**ChIP-qPCR**

Chromatin immunoprecipitation (ChIP) experiments for H3K4me3 were performed as previously described with some modifications[1]. THP-1 cells were cross-linked with 1% formaldehyde in PBS for 10 minutes at room temperature, followed by quenching with 125 mM glycine for an additional 5 minutes. The cells were then lysed in SDS buffer, and chromatin was sheared using a sonicator (Bioruptor) to produce DNA fragments ranging from 200 to 500 bp in size. For immunoprecipitation, 5 µg of anti-H3K4me3 antibody or control IgG was used. After immunoprecipitation, eluted DNA fragments were analyzed using quantitative real-time PCR (qPCR). As an input control, 5% of the sonicated DNA was directly purified prior to immunoprecipitation and evaluated using the same primer set.

##### RNA Sequencing and Data Analysis

Total RNA was extracted from THP1 cells treated with DMSO or 30 nM HHT using Trizol reagent. RNA libraries were constructed and sequenced by Sangon Biotech (Shanghai). Briefly, RNA quality was assessed using a Qubit 2.0 RNA Assay Kit, and libraries were prepared using the Hieff NGSTM MaxUp Dual-mode mRNA Library Prep Kit for Illumina®. Amplified libraries were purified with Hieff NGS™ DNA Selection Beads (0.9×, Beads:DNA =1:1), quantified using the Qubit DNA Assay Kit, and pooled at equimolar ratios. Sequencing was performed on an Illumina HiSeq™ platform to generate 150 bp paired-end reads. Raw reads were quality-checked with FastQC and trimmed using Trimmomatic. High-quality reads were aligned to the reference genome using HISAT2. Read distribution and redundancy were evaluated with RSeQC and Qualimap. Gene expression levels were quantified using StringTie (normalized by TPM). Differential expression analysis was performed with DESeq2 (p-value $< 0.05$), and gene co-expression analysis was conducted using WGCNA. Pathway enrichment analysis for differentially expressed genes was performed using GO, KEGG, and Reactome Gene Sets.

#### QUANTIFICATION AND STATISTICAL ANALYSIS

Data were analyzed using GraphPad Prism 7 and are presented as mean ± SEM. Differences between groups were evaluated using a two-tailed Student's t-test, with a significance threshold set at $p < 0.05$. Survival analysis was conducted using the UALCAN web portal (http://ualcan.path.uab.edu), which facilitates the analysis of TCGA cancer datasets. Differential gene expression was analyzed using limma and DESeq2 within Hiplot Pro (https://hiplot.com.cn/), a comprehensive platform for biomedical data analysis and visualization. Pathway enrichment analysis was performed using GO/KEGG tools in Hiplot Pro and Reactome Gene Sets in Metascape (https://metascape.org/).

**Acknowledgement**

We extend our gratitude to Shanghai Tengyun Biotechnology Co., Ltd. for developing the Hiplot Pro platform (https://hiplot.com.cn/), which provided essential technical support and analytical tools for our research. We also thank the Metascape platform (https://metascape.org/) for offering valuable resources and technical assistance for data analysis and visualization[2]. Additionally, we appreciate the UALCAN team for providing access to their robust analysis tools, which significantly facilitated our survival analysis of TCGA data in this study[3]. We also thank Dr. Zunling Li for providing key experimental materials and for his discussions and assistance throughout the project.

**Table S1. Primers for RT-qPCR, Related to STAR Method.**

| ID | Sequence | Source |
|---|---|---|
| LILRB4-Fwd | **5'**-CATCCATGACAGAGGACTATGC-**3'** | Sangon Biotech, China |
| LILRB4-Rev | **5'**-GGGCTGAAAGGGTGGGTTTA-**3'** | Sangon Biotech, China |
| FTO-Fwd | **5'**-ACTTGGCTCCCTTATCTGACC-**3'** | Sangon Biotech, China |
| FTO-Rev | **5'**-TGTGCAGTGTGAGAAAGGCTT-**3'** | Sangon Biotech, China |
| MLL1-Fwd | **5'**-AAGAGCAGGTAAACTCTCTCCTC-**3'** | Sangon Biotech, China |
| MLL1-Rev | **5'**-TTCCTCTCCGTCGTACAATTTG-**3'** | Sangon Biotech, China |
| cMYC-Fwd | **5'**-GGCTCCTGGCAAAAGGTCA-**3'** | Sangon Biotech, China |
| cMYC-Rev | **5'**-CTGCGTAGTTGTGCTGATGT-**3'** | Sangon Biotech, China |
| RARa-Fwd | **5'**-CCAGCTCCAACAGAAGCAG-**3'** | Sangon Biotech, China |
| RARa-Rev | **5'**-AGGCCTCTGTCCAAGGAGTC-**3'** | Sangon Biotech, China |
| GAPDH-Fwd | **5'**-AGCAAGAGCACAAGAGGAAG-**3'** | Sangon Biotech, China |
| GAPDH-Rev | **5'**-GGTTGAGCACAGGGTACTTT-**3'** | Sangon Biotech, China |

**Table S2. Oligonucleotides for shRNA Construction, Related to STAR Methods.**

| ID | Sequence | Source |
|---|---|---|
| Sh FTO-1 | **5'**-TCACCAAGGAGACTGCTATTT-**3'** | Sangon Biotech, China |
| Sh FTO-2 | **5'**-CGGTTCACAACCTCGGTTTAG-**3'** | Sangon Biotech, China |
| Sh NC | **5'**-TTCTCCGAACGTGTCACGT-**3'** | Sangon Biotech, China |
| Sh GSK-3β | **5'**-ACTAGAGGGCAGAGGTAAAT-**3'** | Sangon Biotech, China |
| Sh MLL-1 | **5'**-GCACTGTTAAACATTCCACTT-**3'** | Sangon Biotech, China |
| Sh MLL-2 | **5'**-GATTATGACCCTCCAATTAAA-**3'** | Sangon Biotech, China |

**Figure Legends:**

**Figure S1: LILRB4 Expression is Closely Associated with Clinical Patient Survival Rates**

(A) Box plot illustrating the expression levels of LILRB4 in various subtypes of AML patients, as analyzed using online tools from the TCGA database. The data indicate a specifically elevated expression of LILRB4 in AML-M4/M5 patients.

(B) Kaplan-Meier survival curve generated from TCGA database analysis showing that patients with high LILRB4 expression exhibit lower survival rates. The curve compares two groups: one with high LILRB4 expression (n=43) represented in red, and the other with low LILRB4 expression (n=120) represented in blue. The log-rank test p-value is indicated as p=0.042, demonstrating a statistically significant difference in survival between the two groups. These results suggest that LILRB4 expression levels may serve as a prognostic indicator in the disease under study, with higher expression correlating with poorer outcomes.

**Figure S2: Analysis of H3K4me3 Enrichment at LILRB4 Locus in AML Cell Lines**

(A) ChIP-seq data for H3K4me3 in AML cell lines obtained from the Cistrome Data Browser, demonstrating a significant enrichment of H3K4me3 levels at the LILRB4 promoter region in cell lines with high LILRB4 expression. The histograms represent the normalized read counts of H3K4me3 marks across the genome, with the bottom track showing the genomic location of LILRB4, highlighting the regions of interest. The data suggest that high LILRB4-expressing cells exhibit increased H3K4me3 histone methylation at the LILRB4 locus, potentially influencing gene expression and regulation.

**Figure S3: HHT Modulates the Expression of Epigenetic Regulators in MV4-11 Cells**

(A) Western blot analysis of various epigenetic complexes in MV4-11 cells after treatment with 0-160 nM HHT for 48 hours. The expression levels of the m6A demethylase FTO, and components of the H3K4me3/H3K79me2 complex (MENIN, MLL-N, MLL-C), H3K27me3 complex, and PRMTs were assessed. The results show a significant downregulation in the expression of FTO, MENIN, MLL-N, and MLL-C upon HHT treatment. The red plus signs indicate the bands of interest. This suggests that HHT may exert its effects on AML cells by modulating the expression of key epigenetic regulatory factors.

**Figure S4: FTO Regulates LILRB4 Expression Levels**

(A-B) Total RNA was extracted from MV4-11 cells, and RT-qPCR analysis revealed a decrease in LILRB4 mRNA expression upon FTO knockdown.

(C) Western blot analysis of MV4-11 cells with FTO knockdown showed a suppression of LILRB4 protein levels.

(D) Treatment of NOMO-1 cells with the FTO enzyme inhibitor FB23-2 led to a reduction in LILRB4 expression. These results indicate that FTO plays a role in controlling the expression of LILRB4 at both the transcriptional and translational levels.

**Figure S5: HHT Promotes FTO Degradation via Ubiquitination to Suppress FTO Protein Expression**

(A) RNA-seq analysis indicates an increase in FTO transcript levels upon HHT treatment, suggesting that HHT regulation of FTO does not occur at the transcriptional level.

(B) Co-immunoprecipitation (co-IP) assay of THP1 cell lysates treated with DMSO and 50 nM HHT for 36 hours, demonstrating that HHT inhibits FTO protein expression by promoting its ubiquitination and subsequent degradation.

(C) Image of purified FTO protein in vitro.

(D) Enzyme activity assay of purified FTO protein treated with increasing concentrations of HHT, showing that HHT treatment does not affect the enzymatic activity of FTO. This figure collectively suggests that HHT suppresses FTO protein expression by enhancing its ubiquitination and degradation rather than through transcriptional regulation.

**Figure S6: HHT Diminishes the Inhibitory Effect of AML Cells on T Cell Viability**

(A) Representative FACS plots relative to Fig. 6D are shown. Flow cytometry analysis depicting the co-culture of T cells with THP1 cells that were pre-treated with either 60 nM HHT or an equivalent volume of DMSO. After treatment, cells were washed to eliminate residual compounds prior to co-culturing with T cells at a 1:1 ratio for 16 to 20 hours. The DMSO control group displays a marked decrease in the the proportion of T cells. Conversely, the group treated with 60 nM HHT exhibits a heightened percentage of cytotoxic T cells. This suggests that HHT treatment substantially lessens the suppressive influence of THP1 cells on T cell proliferation, which in turn fosters the restoration of T cell-mediated immune responses.

**Figure S7: HHT Inhibits Immunevasion in Humanized Mouse AML Xenografts**

(A) Relative to Fig.7 A. Tumors were isolated and imaged as shown. Comparative tumor size graph showing that treatment with HHT significantly reduces tumor size in the $THP^{WT}$ group, whereas the $LILRB4^{KO}$ group shows no difference in tumor growth regardless of HHT treatment.

(B) Quantification of tumor sizes for each group, providing a numerical representation of the tumor volume changes in response to treatment.

(C) Quantification of tumor weights for each group, illustrating the impact of HHT treatment on tumor mass.

(D) The FACS histogram of LILRB4 expression of tumor cells. Analysis of LILRB4 expression in tumor cells, indicating that HHT treatment decrease LILRB4 expression in the THP1 ($LILRB4^{WT}$) group.

(E-I) Relative to Fig.7.G-K. The FACS plots of expression of CD3, CD8, CD4, and Ki-67 markers in tumor-infiltrating lymphocytes analyzed by FACS. In the $LILRB4^{WT}$ group, HHT treatment significantly affects various immune parameters. In contrast, in the $LILRB4^{KO}$ group, there is no difference in immune marker expression with or without HHT treatment, suggesting that the inhibitory effect of HHT on tumor immune evasion is dependent on the presence of LILRB4.

**Figure S8: Gating strategies used in FACs analysis for human PBMC reconstruction assay.**

Gating strategy to identify transfer human PBMC in vivo after 7 days transplant such as Fig. 7 and Supplementary Fig.7.

### Supplementary Figure 1. (Huang et al.)

A.

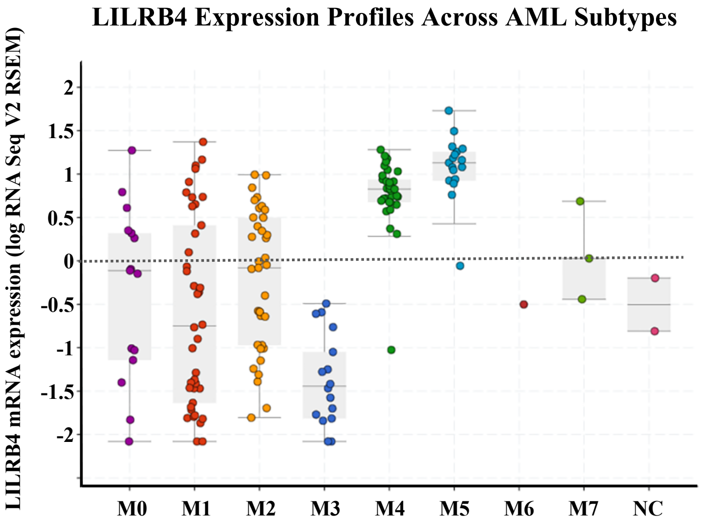


B.

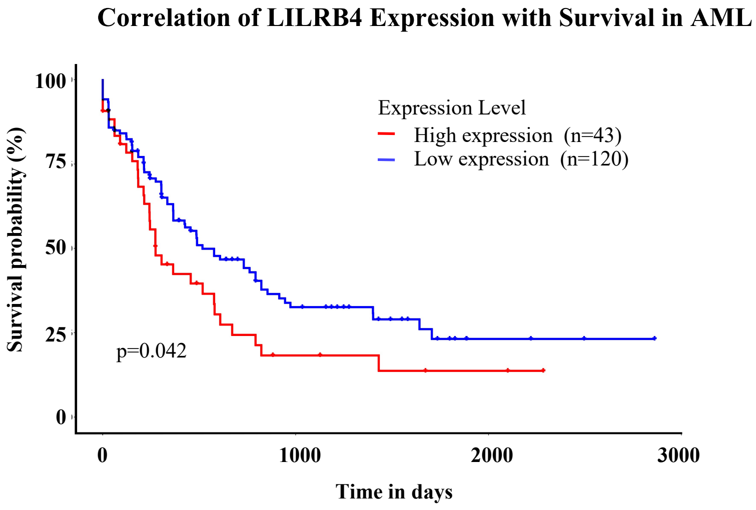

**Supplementary Figure 2. (Huang et al.)**

**A.**

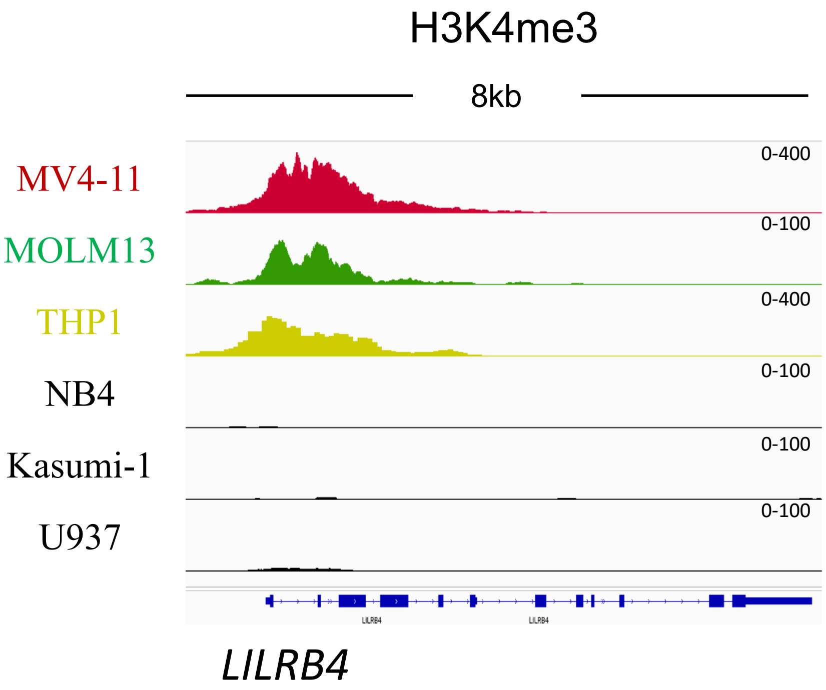

Supplementary Figure 3. (Huang et al.)

A.

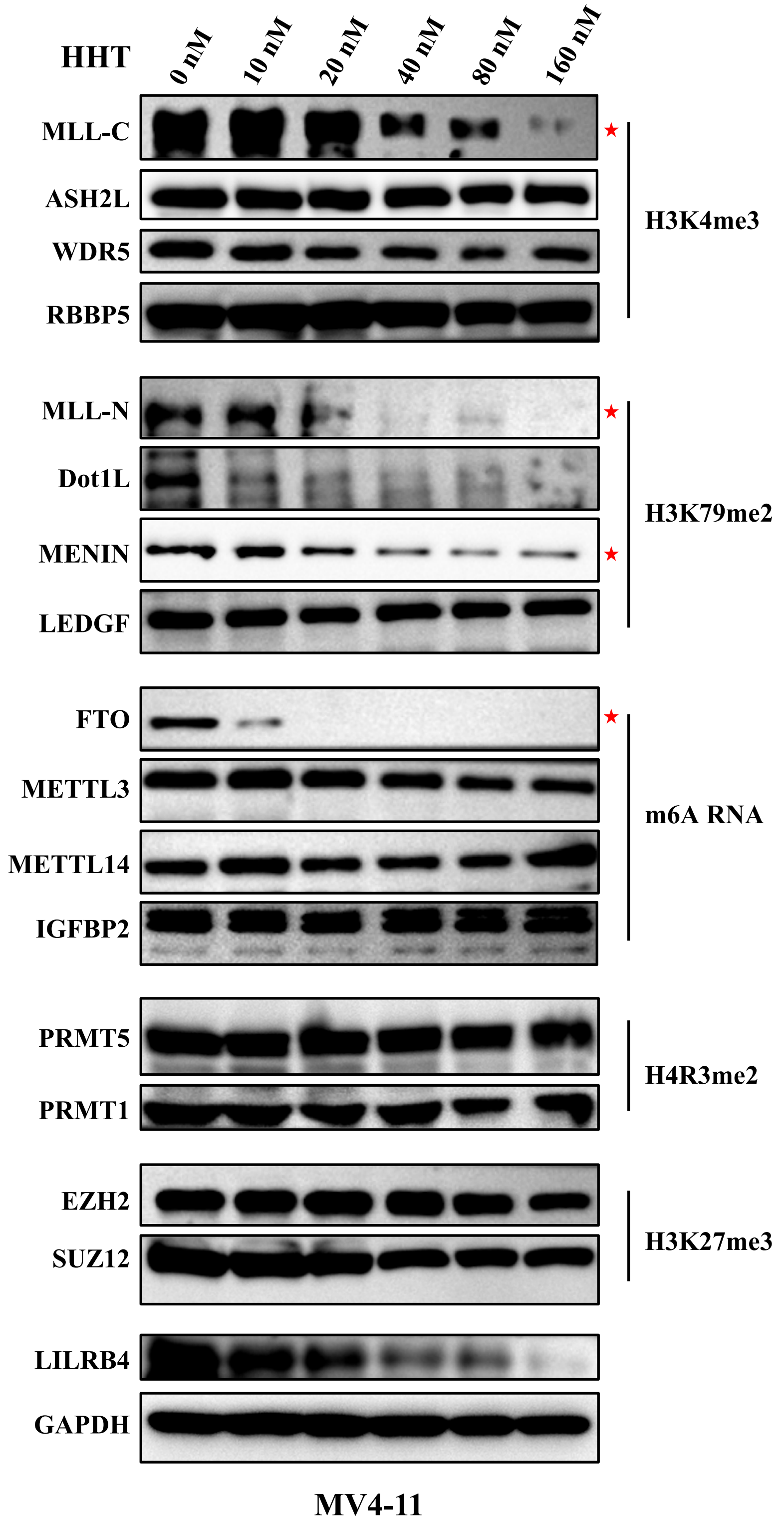

### Supplementary Figure 4. (Huang et al.)

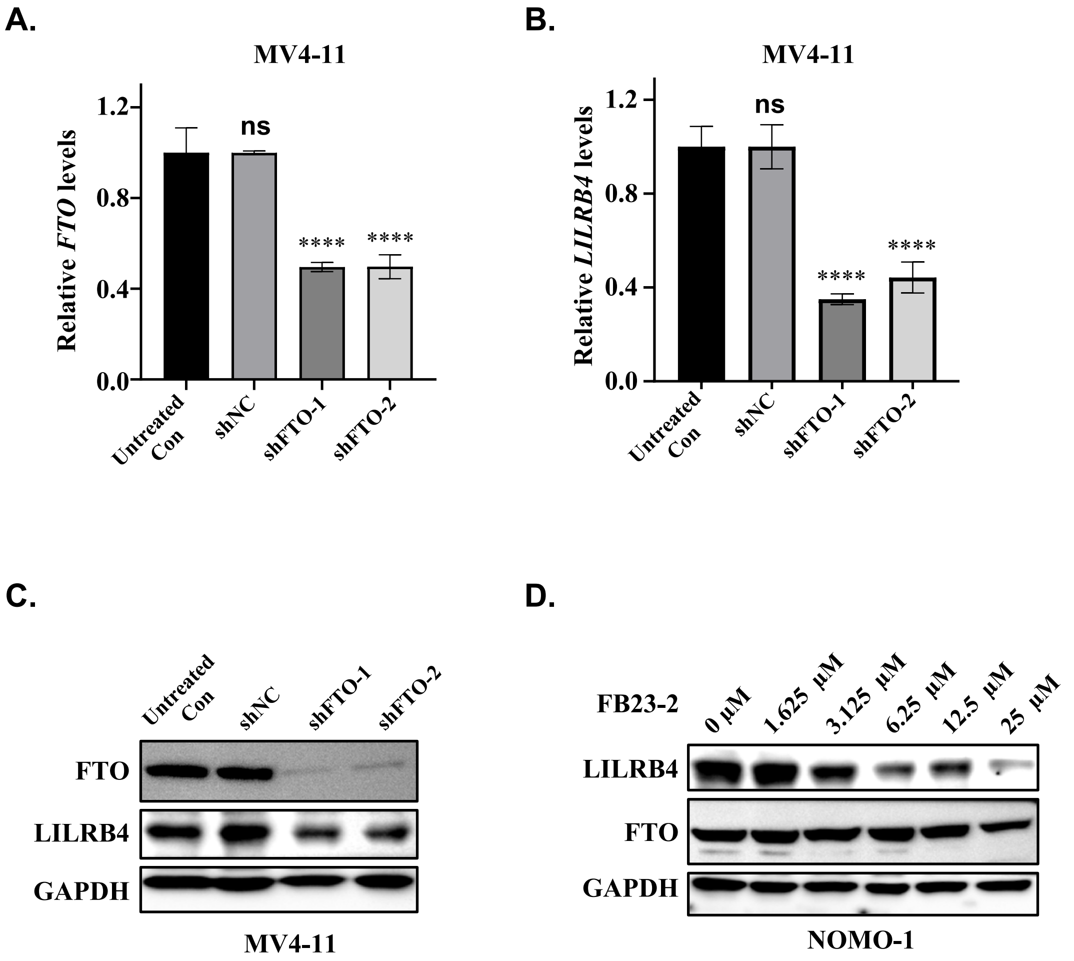

#### Supplementary Figure 5. (Huang et al.)

**A.**

| log2FoldChange | pValue | qValue | result | GeneName |
|---|---|---|---|---|
| 2.06765014 | 0.006192058 | 0.014628001 | up | FTO |

**B.**

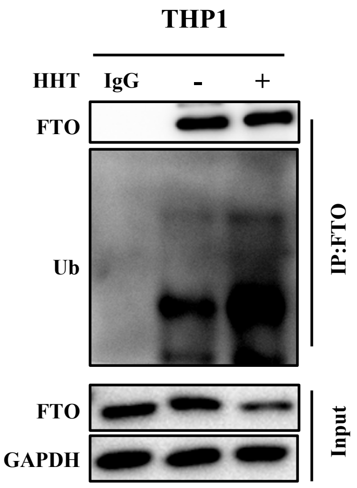


**C.**

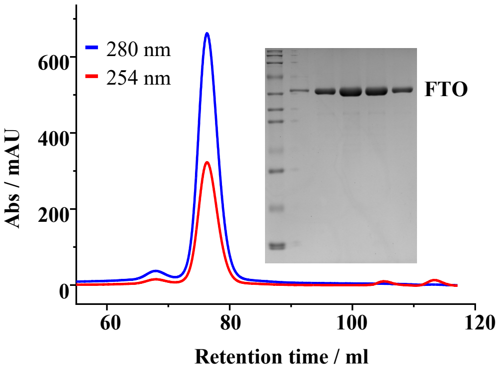


**D.**

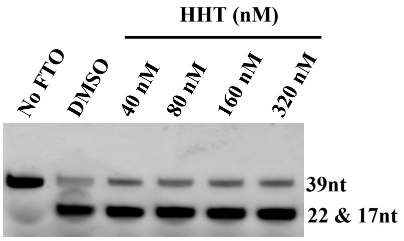

#### Supplementary Figure 6. (Huang et al.)

**A.**

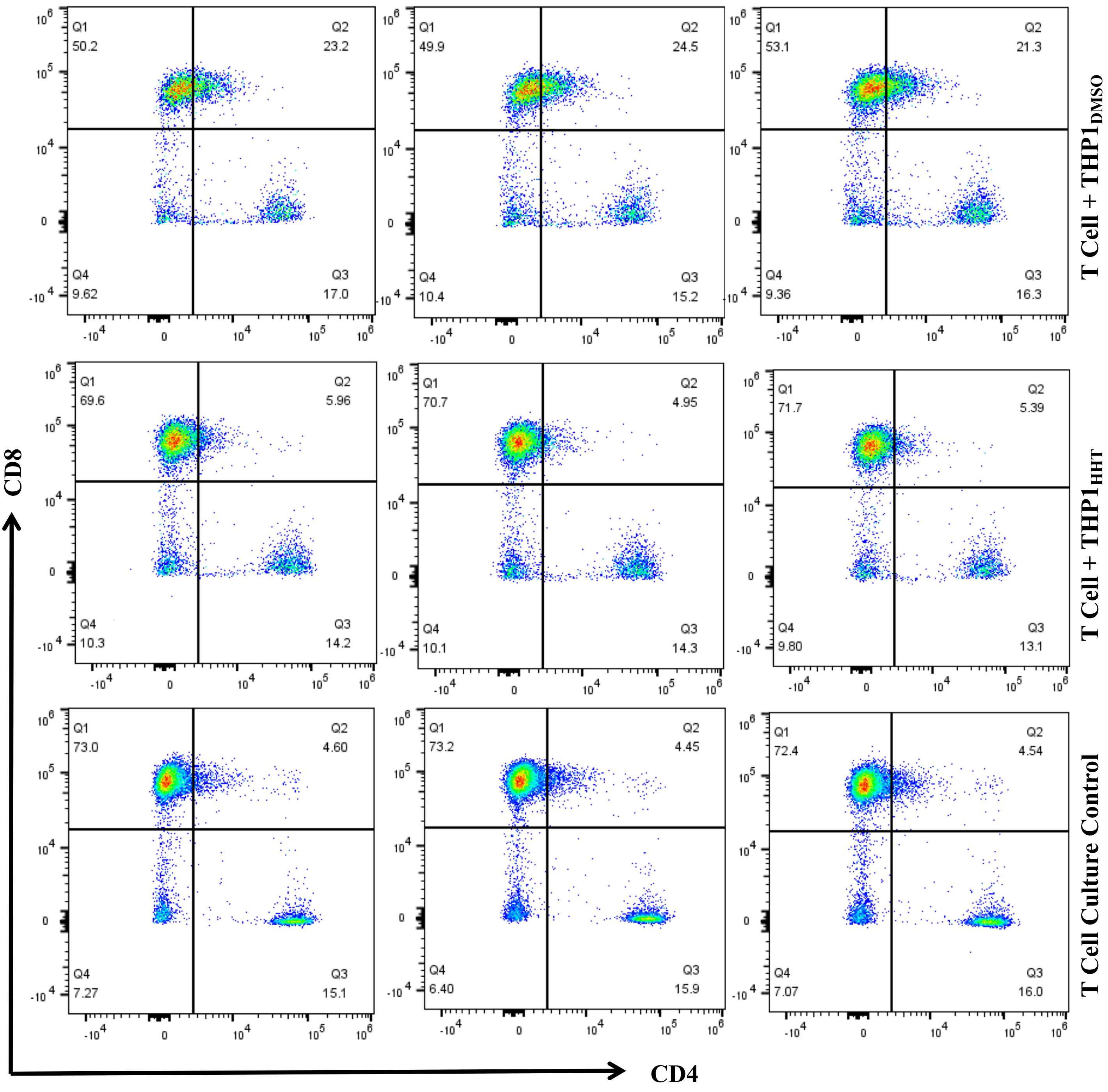

#### Supplementary Figure 7. (Huang et al.)

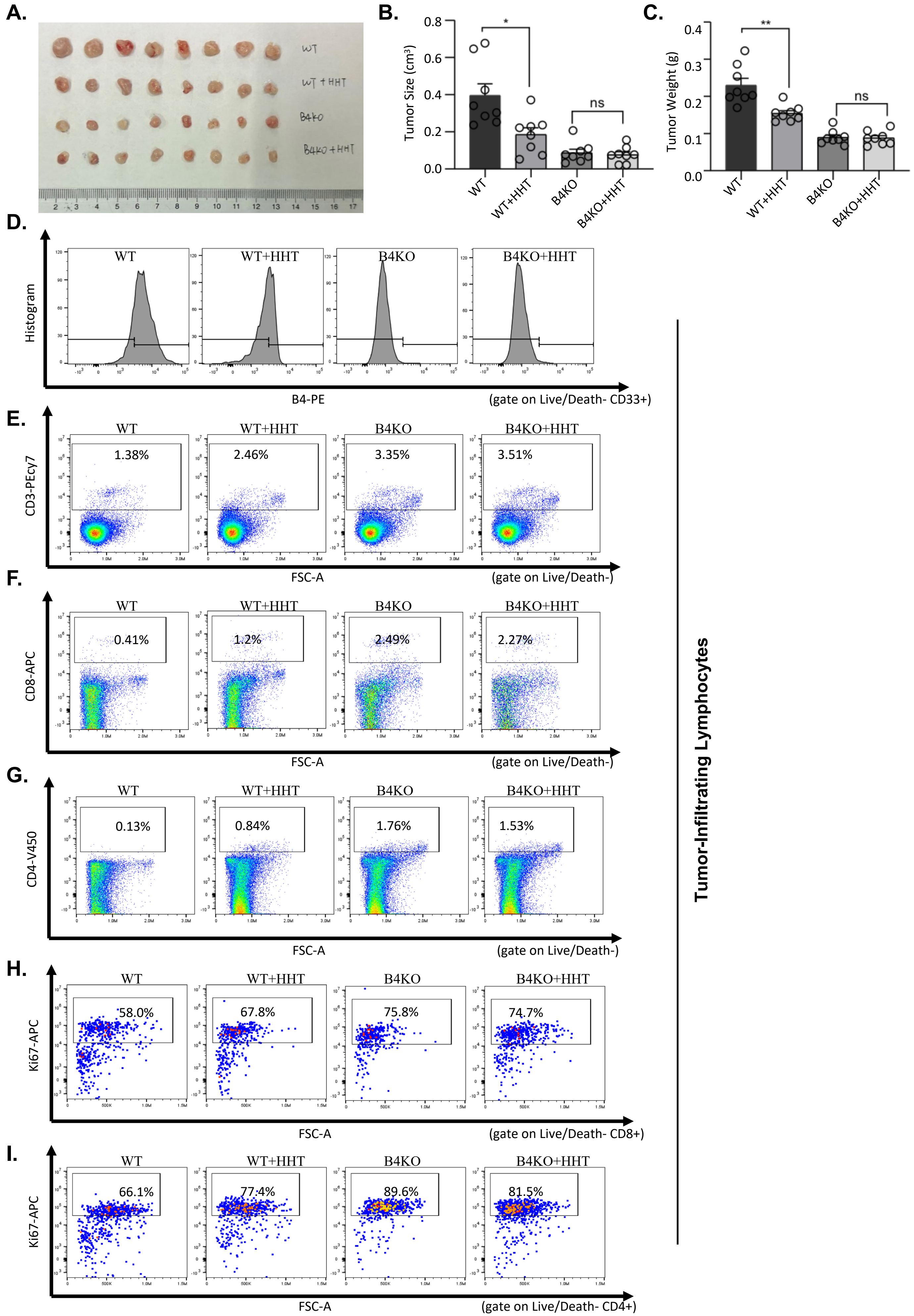

### Supplementary Figure 7. (Huang et al.)

## J.

Human PBMC reconstruction assay (7days after transplant)

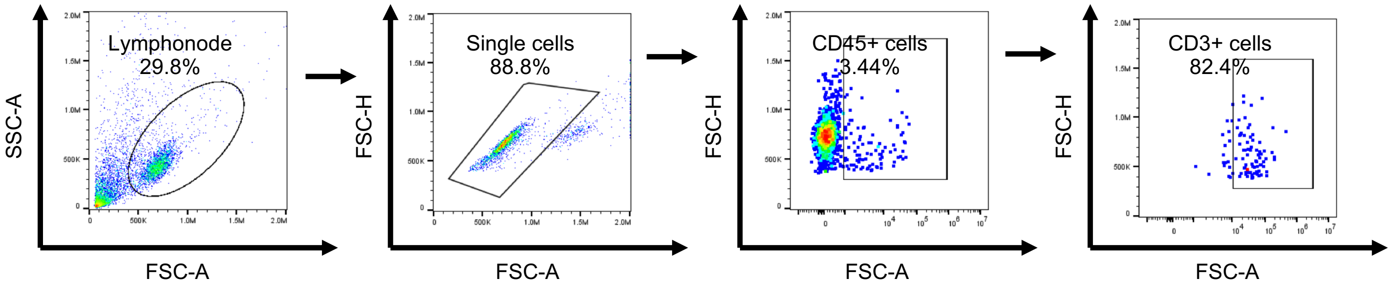
